# Characterization of Influenza A HA and NA Subtypes Using Protein Sequences

**DOI:** 10.64898/2026.07.22.740123

**Authors:** Yunfei Wang, Fabiola Vacca, Wes Rountree, Kevin Wiehe, Max M. He, M. Anthony Moody

## Abstract

**Motivation:** Influenza is an infectious disease associated with excess human deaths. For influenza A virus (IAV), the surface antigens hemagglutinin (HA) and neuraminidase (NA) define the specific subtype. The high mutation rate of IAV can make clinical testing and assigning subtype after sequencing challenging. Accurate subtype classification is important for tracking circulating IAV strains that may impact human health.

**Results:** We analyzed a large IAV protein sequence dataset with known HA and NA subtypes. Using logistic regression and random forest approaches, we identified a small set of subtype-associated amino acids, which were then used to develop a subtype characterization method with near-optimal accuracy. Further analysis indicated that signal sequence peptide variation and indels in HA and NA explained the unique combination of subtype-associated amino acids.

## Introduction

Influenza is a contagious respiratory disease caused by influenza viruses. Humans can be infected with influenza A, B, and C, and influenza A virus (IAV) and influenza B cause seasonal outbreaks leading to severe illness or death (Uyeki et al. 2022). Seasonal influenza has significant global health and economic impacts, causing between 291,000 to 645,000 deaths annually worldwide (Venkatramanan et al., 2021). The World Health Organization (WHO) estimates that seasonal influenza epidemics result in 3 to 5 million cases of severe illness (hospitalizations) annually worldwide (Paget et al. 2024). Revisiting influenza-hospitalization estimates from the Burden of Influenza and Respiratory Syncytial Virus Disease (BIRD) project using different extrapolation methods (Paget et al. 2024). Since the early 1900s, four flu pandemics have occurred: the 1918 H1N1 Spanish flu, the 1957 H2N2 Asian flu, the 1968 H3N2 Hong Kong flu, and the 2009 H1N1 swine flu, with the 1918 pandemic being the deadliest, causing 50 million deaths worldwide (Dowdle et al., 1999).

The influenza virus genome is ∼13kb and encodes 13 proteins: hemagglutinin (HA), neuraminidase (NA), M1 matrix protein (M1), M2 ion channel protein (M2), nuclear protein (NP), nonstructural proteins (NS1, NS2), and the RNA polymerase complex (PB1, PB2, PA). IAVs are currently divided into 18 HA subtypes and 11 NA subtypes based on their surface glycoproteins, while influenza B and C viruses are less genetically diverse and are not usually further subdivided (Shao et al., 2017). Influenza D viruses affecting cattle with spillovers to other animals are not known to cause illness in people, hence are of less concern (CDC, n.d.).

People can be infected by influenza repeatedly due to the ongoing evolution of these viruses, which helps them evade immunity gained from past infections or vaccinations and facilitates their transmission between people. At present, the most common IAV subtypes infecting humans are H1N1 and H3N2, but sporadic infections with other subtypes occur regularly. Accurate classification of IAV subtypes is crucial to surveillance and monitoring efforts to permit effective planning of disease prevention measures.

Methods for characterizing IAV HA and NA subtypes include Hemagglutination Inhibition (HI) assay, Neuraminidase Inhibition (NI) assay, genomic sequencing, immunostaining, antigenic characterization, and mass spectrometry (Hong et al., 2018, LeBlanc et al., 2020, Humayun et al., 2021, Johnson et al., 2012, Schwahn et al., 2009). It is well established that the HI and NI assays have historically been the standard method for HA subtype identification, based on the ability of subtype-specific antibodies to inhibit hemagglutination and Neuraminidase (Lee et al., 2006; Pedersen, 2008; WHO, 2011.). However, HI and NI assays can be cumbersome, requiring well-characterized antigens and antisera for all subtypes. In addition, there is often a need for multiple reagents for each subtype, and interpreting results can be challenging, especially for variants within a subtype or mixed infections. Genomic sequencing, antigenic characterization, and mass spectrometry provide alternative methods for characterizing IAV subtypes, often used in combination with other methods of characterization (Hong et al., 2018), but improved algorithms could help surveillance efforts.

In this study, we developed a simple and highly accurate approach for classifying IAV strains (HA and NA types) by identifying a small set of subtype-associated amino acids that we developed into a simple subtype classification method. To understand why these amino acids were predictive, we investigated the sequences and structural locations of four subtype-predictive reference residues in HA and NA.

## Results

### A combination of four amino acid positions can be used to predict HA and NA subtype

There are 18 HA types, with the proportion of each type shown in Figure 1. The types most commonly present in the database are H1, H3, H9, H5, H7, H4, H6, H10, H11, H2, and H13. The mixed type accounts for 2%, while other minor HA type (H99), including H8, H12, and H14 ∼ H18, make up 1%. For prediction model fitting, a sample size of n=83,872 was used, excluding the mixed type from the total sample size of n=85,645.

**Fig. 1.**
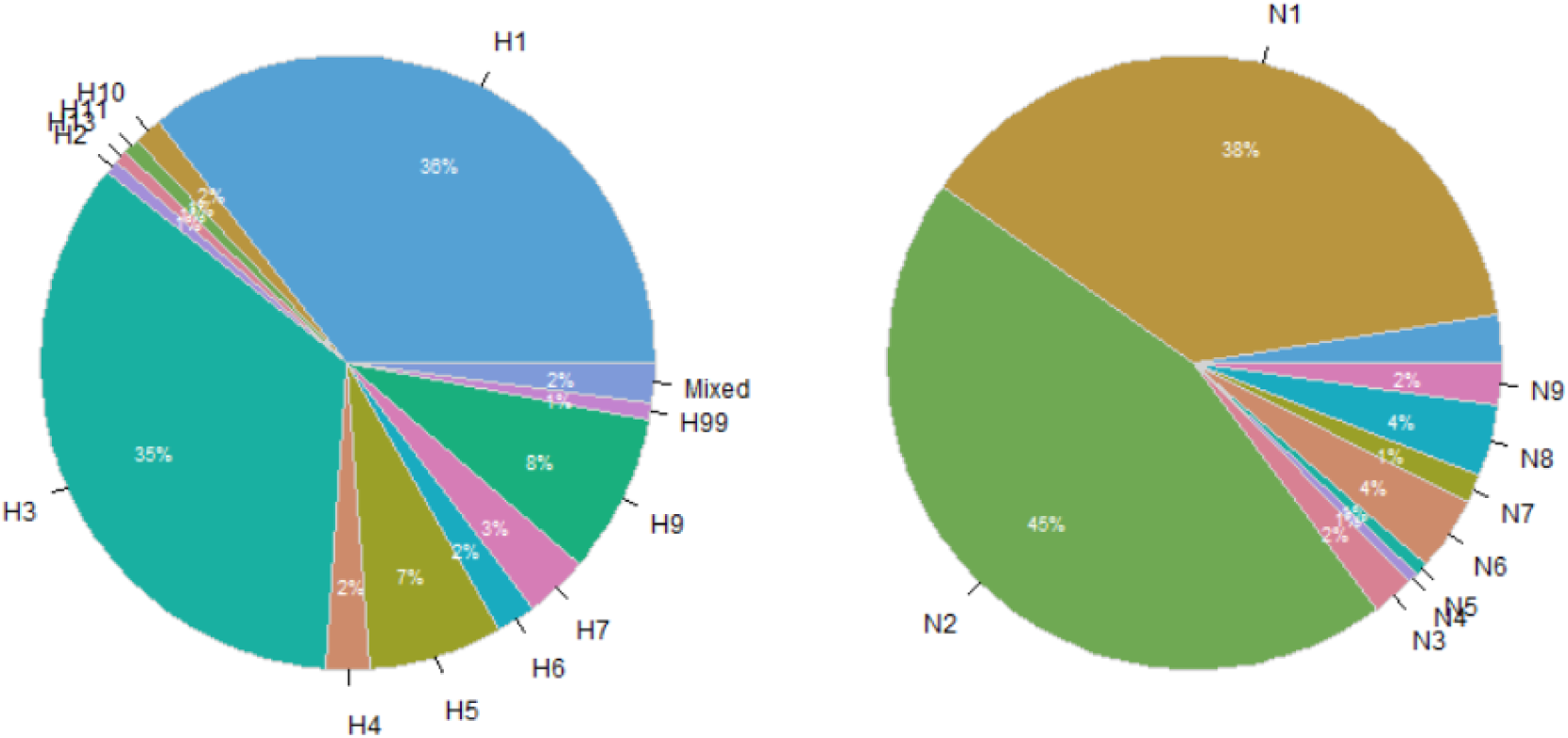
Pie charts for the HA and NA type distribution used in the study (Year 1902 ∼2019).

There are 11 NA types, with the proportion of each type also shown in Figure 1. The types most commonly present are N2, N1, N6, N8, N3, N9, N7, N5 and N4. After excluding 1886 missing NA-type cases and 4 other cases (n=3 for N10, n=1 for N11), a total sample size of n=72,407 was retained for model training and testing.

To classify HA and NA subtypes, we used a univariate logistic regression model to test the association of each amino acid in a different position with the HA or NA subtype. The prediction accuracy for each amino acid was used to select predictors for model fitting. The four predictors with the highest accuracy were used for the final prediction models implemented in the app. Table 1 shows that each major HA subtype is associated with a unique amino acid at Position 125 (Pos125), except H13, which shares the same amino acid G with H7. Similarly, the distributions of the other 7 amino acids at Pos56, Pos45, and Pos48 from HA, Pos111, Pos409, Pos34, and Pos45 from NA were shown to be strongly associated with either the HA or NA subtype shown in Table 1s ∼ Table 7s in the supplemental material. The positions of these amino acids are the perfect candidates for model prediction and are hereafter referenced as subtype-predictive.

**Table 1.**
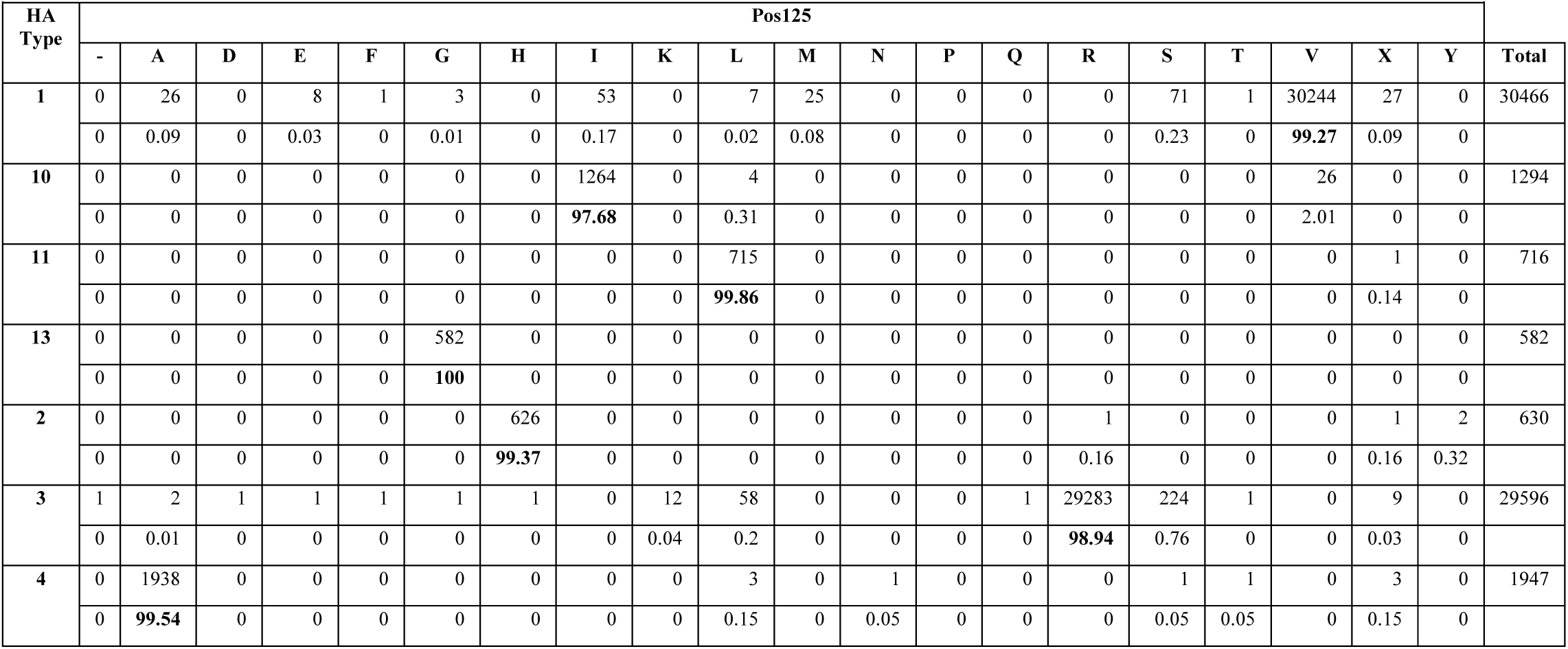
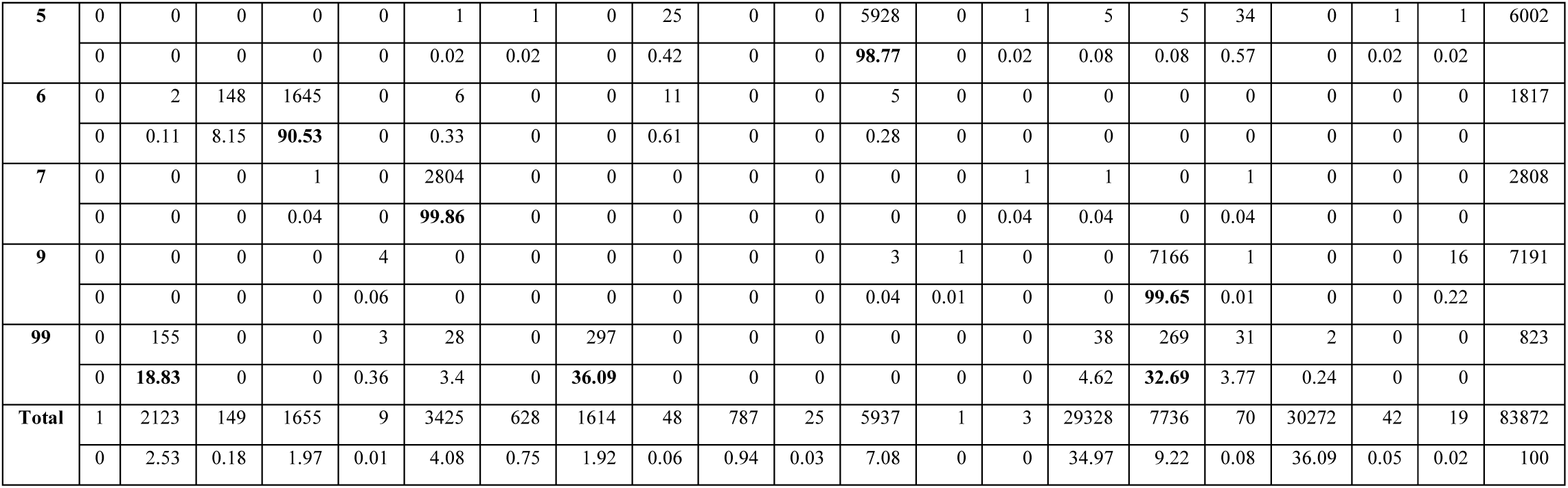
The distribution of amino acids in HA Position 125 among different HA types.

|  |  |  |  |  |  |  |  |  |  |  |  |  |  |  |  |  |  |  |  |  |  |
| --- | --- | --- | --- | --- | --- | --- | --- | --- | --- | --- | --- | --- | --- | --- | --- | --- | --- | --- | --- | --- | --- |
| HA Type | Pos125 |  |  |  |  |  |  |  |  |  |  |  |  |  |  |  |  |  |  |  | Total |
|  | - | A | D | E | F | G | H | I | K | L | M | N | P | Q | R | S | T | V | X | Y |  |
| 1 | 0 | 26 | 0 | 8 | 1 | 3 | 0 | 53 | 0 | 7 | 25 | 0 | 0 | 0 | 0 | 71 | 1 | 30244 | 27 | 0 | 30466 |
|  | 0 | 0.09 | 0 | 0.03 | 0 | 0.01 | 0 | 0.17 | 0 | 0.02 | 0.08 | 0 | 0 | 0 | 0 | 0.23 | 0 | 99.27 | 0.09 | 0 |  |
| 10 | 0 | 0 | 0 | 0 | 0 | 0 | 0 | 1264 | 0 | 4 | 0 | 0 | 0 | 0 | 0 | 0 | 0 | 26 | 0 | 0 | 1294 |
|  | 0 | 0 | 0 | 0 | 0 | 0 | 0 | 97.68 | 0 | 0.31 | 0 | 0 | 0 | 0 | 0 | 0 | 0 | 2.01 | 0 | 0 |  |
| 11 | 0 | 0 | 0 | 0 | 0 | 0 | 0 | 0 | 0 | 715 | 0 | 0 | 0 | 0 | 0 | 0 | 0 | 0 | 1 | 0 | 716 |
|  | 0 | 0 | 0 | 0 | 0 | 0 | 0 | 0 | 0 | 99.86 | 0 | 0 | 0 | 0 | 0 | 0 | 0 | 0 | 0.14 | 0 |  |
| 13 | 0 | 0 | 0 | 0 | 0 | 582 | 0 | 0 | 0 | 0 | 0 | 0 | 0 | 0 | 0 | 0 | 0 | 0 | 0 | 0 | 582 |
|  | 0 | 0 | 0 | 0 | 0 | 100 | 0 | 0 | 0 | 0 | 0 | 0 | 0 | 0 | 0 | 0 | 0 | 0 | 0 | 0 |  |
| 2 | 0 | 0 | 0 | 0 | 0 | 0 | 626 | 0 | 0 | 0 | 0 | 0 | 0 | 0 | 1 | 0 | 0 | 0 | 1 | 2 | 630 |
|  | 0 | 0 | 0 | 0 | 0 | 0 | 99.37 | 0 | 0 | 0 | 0 | 0 | 0 | 0 | 0.16 | 0 | 0 | 0 | 0.16 | 0.32 |  |
| 3 | 1 | 2 | 1 | 1 | 1 | 1 | 1 | 0 | 12 | 58 | 0 | 0 | 0 | 1 | 29283 | 224 | 1 | 0 | 9 | 0 | 29596 |
|  | 0 | 0.01 | 0 | 0 | 0 | 0 | 0 | 0 | 0.04 | 0.2 | 0 | 0 | 0 | 0 | 98.94 | 0.76 | 0 | 0 | 0.03 | 0 |  |
| 4 | 0 | 1938 | 0 | 0 | 0 | 0 | 0 | 0 | 0 | 3 | 0 | 1 | 0 | 0 | 0 | 1 | 1 | 0 | 3 | 0 | 1947 |
|  | 0 | 99.54 | 0 | 0 | 0 | 0 | 0 | 0 | 0 | 0.15 | 0 | 0.05 | 0 | 0 | 0 | 0.05 | 0.05 | 0 | 0.15 | 0 |  |
| 5 | 0 | 0 | 0 | 0 | 0 | 1 | 1 | 0 | 25 | 0 | 0 | 5928 | 0 | 1 | 5 | 5 | 34 | 0 | 1 | 1 | 6002 |
|  | 0 | 0 | 0 | 0 | 0 | 0.02 | 0.02 | 0 | 0.42 | 0 | 0 | 98.77 | 0 | 0.02 | 0.08 | 0.08 | 0.57 | 0 | 0.02 | 0.02 |  |
| 6 | 0 | 2 | 148 | 1645 | 0 | 6 | 0 | 0 | 11 | 0 | 0 | 5 | 0 | 0 | 0 | 0 | 0 | 0 | 0 | 0 | 1817 |
|  | 0 | 0.11 | 8.15 | 90.53 | 0 | 0.33 | 0 | 0 | 0.61 | 0 | 0 | 0.28 | 0 | 0 | 0 | 0 | 0 | 0 | 0 | 0 |  |
| 7 | 0 | 0 | 0 | 1 | 0 | 2804 | 0 | 0 | 0 | 0 | 0 | 0 | 0 | 1 | 1 | 0 | 1 | 0 | 0 | 0 | 2808 |
|  | 0 | 0 | 0 | 0.04 | 0 | 99.86 | 0 | 0 | 0 | 0 | 0 | 0 | 0 | 0.04 | 0.04 | 0 | 0.04 | 0 | 0 | 0 |  |
| 9 | 0 | 0 | 0 | 0 | 4 | 0 | 0 | 0 | 0 | 0 | 0 | 3 | 1 | 0 | 0 | 7166 | 1 | 0 | 0 | 16 | 7191 |
|  | 0 | 0 | 0 | 0 | 0.06 | 0 | 0 | 0 | 0 | 0 | 0 | 0.04 | 0.01 | 0 | 0 | 99.65 | 0.01 | 0 | 0 | 0.22 |  |
| 99 | 0 | 155 | 0 | 0 | 3 | 28 | 0 | 297 | 0 | 0 | 0 | 0 | 0 | 0 | 38 | 269 | 31 | 2 | 0 | 0 | 823 |
|  | 0 | 18.83 | 0 | 0 | 0.36 | 3.4 | 0 | 36.09 | 0 | 0 | 0 | 0 | 0 | 0 | 4.62 | 32.69 | 3.77 | 0.24 | 0 | 0 |  |
| Total | 1 | 2123 | 149 | 1655 | 9 | 3425 | 628 | 1614 | 48 | 787 | 25 | 5937 | 1 | 3 | 29328 | 7736 | 70 | 30272 | 42 | 19 | 83872 |
|  | 0 | 2.53 | 0.18 | 1.97 | 0.01 | 4.08 | 0.75 | 1.92 | 0.06 | 0.94 | 0.03 | 7.08 | 0 | 0 | 34.97 | 9.22 | 0.08 | 36.09 | 0.05 | 0.02 | 100 |

We evaluated the models using accuracy, precision, recall, and F1 score. The logistic regression model outperformed the random forest model (random forest model results not shown), although the logistic regression occasionally crashed when an amino acid in the test dataset did not exist in the training dataset. To address this issue, we used the random forest for predictions in such cases.

As can be seen in Table 2, the accuracy of using a single predictor (the amino acid at Position 125 (Pos125)) is 0.976 with a 95% CI of (0.974, 0.978). To enhance prediction accuracy, four predictors (Pos125, Pos56, Pos45, and Pos48) were selected to build the final prediction model in characterizing HA types. Similarly, the accuracy of a single predictor model using variable Pos111 is 0.972 with a 95% CI of (0.969, 0.974). Four predictors (Pos111, Pos409, Pos34, and Pos45) were chosen for the final model in characterizing NA types.

**Table 2.** Accuracy for HA or NA logistic regression models.

|  | Type | Predictors | Estimate | 95%<br>lower<br>bound | 95%<br>upper<br>bound |
| --- | --- | --- | --- | --- | --- |
| Simple Model | HA | Pos125 | 0.9763 | 0.9742 | 0.9784 |
| Simple Model | NA | Pos111 | 0.9718 | 0.9693 | 0.9743 |
| Final Model | HA | Pos125, Pos56, Pos45 and Pos48 | 0.9991 | 0.9987 | 0.9995 |
| Final Model | NA | Pos111, Pos409, Pos34, and Pos45 | 0.9994 | 0.9990 | 0.9998 |

The accuracy in predicting HA type using the final logistic regression model was 0.9991 with a 95% CI of (0.9987, 0.9995), and for NA type, it was 0.9994 with a 95% CI of (0.9990, 0.9998) shown in Table 2. Figures 2 and 3 show metrics of PREC, REC, and F1 score. Scores ranged from 0.992 to 1 for HA types except for H99 HA type (REC=0.968, F1 scores=0.983), and 0.995 to 1 for NA types. These final prediction models showed almost perfect accuracy, precision, recall, and F1 scores. Furthermore, the additional independent validation results showed that the HA prediction model has an accuracy of 0.9989 whereas the NA prediction model has an accuracy of 0.9983. Overall, our method shows almost perfect accuracy, precision, recall and F1 score.

**Fig. 2.**
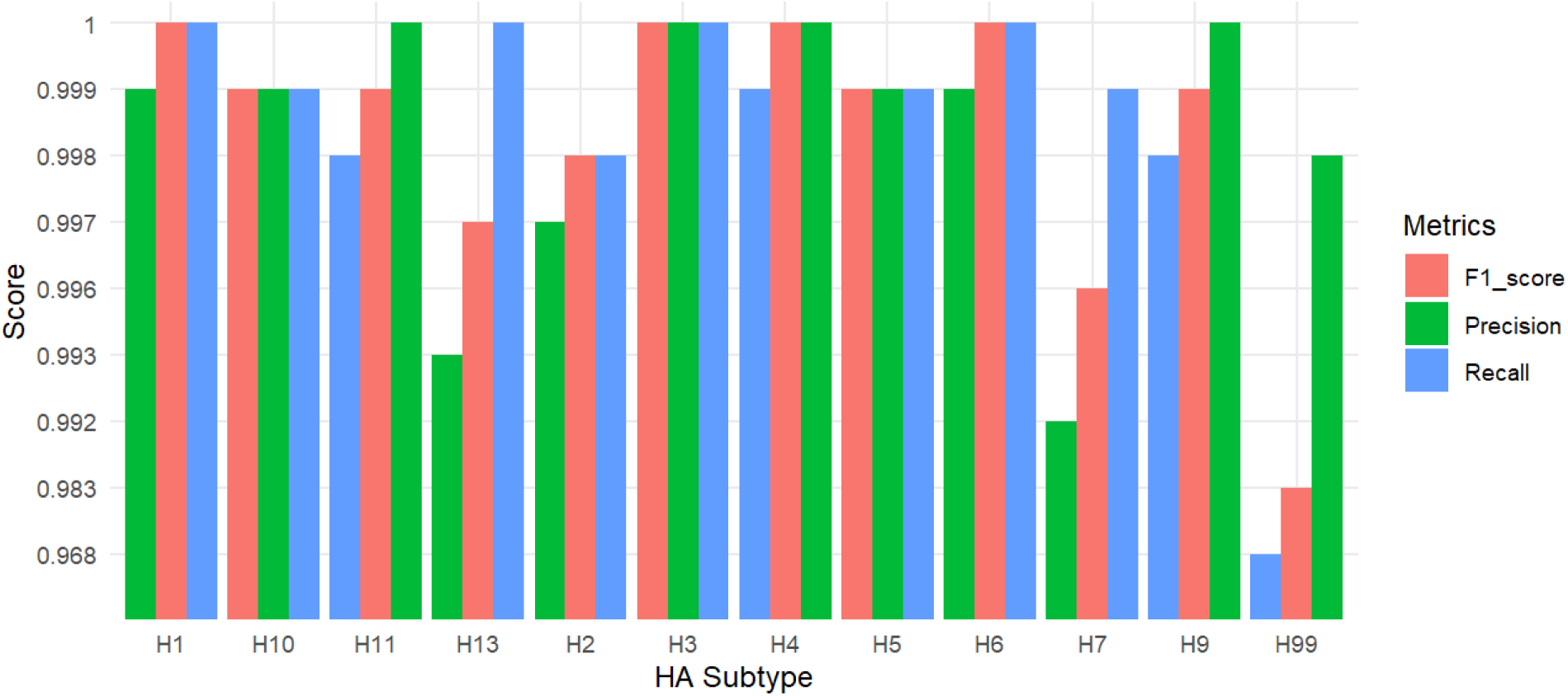
Precision, Recall and F1_score of final logistic regression model by HA Subtype.

**Fig. 3.**
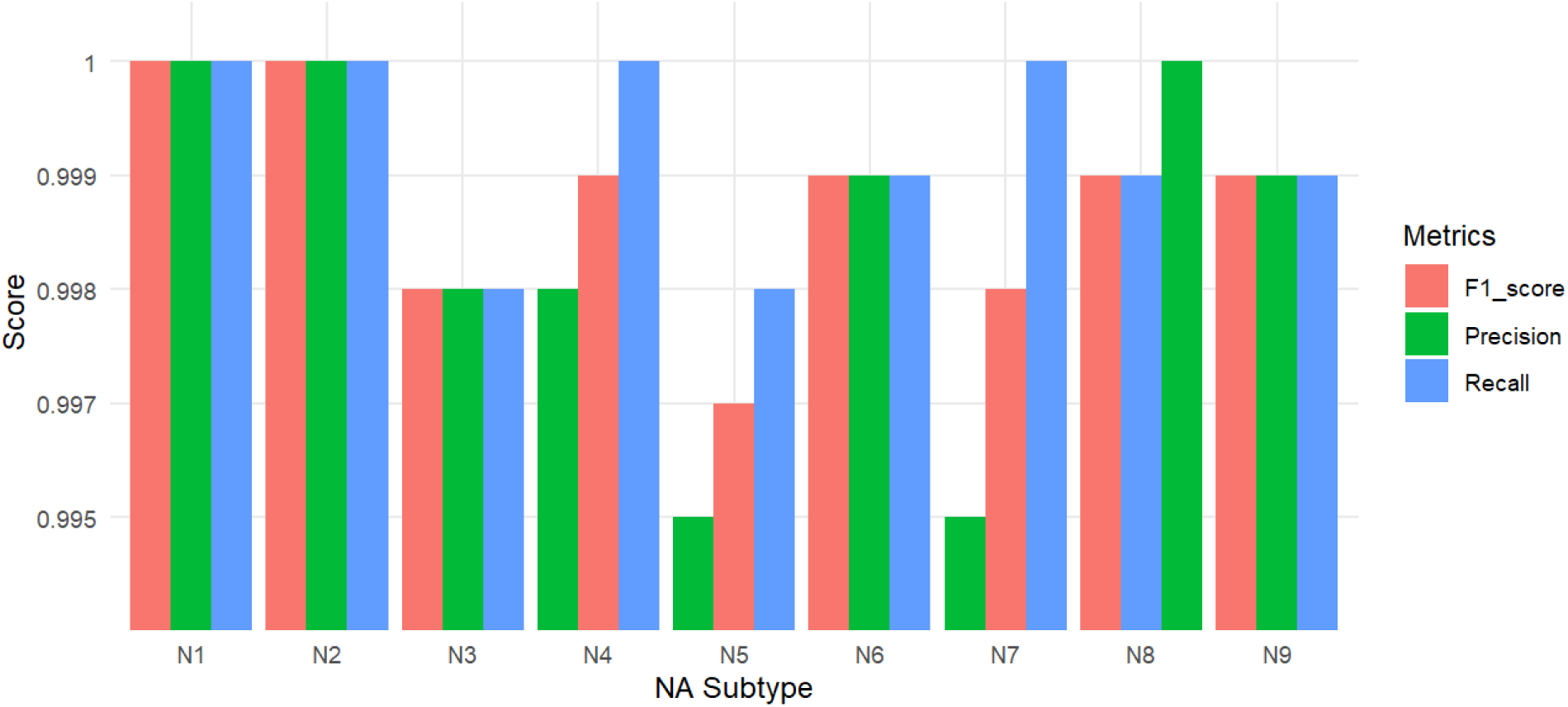
Precision, Recall and F1_score of final logistic regression model by NA Subtype.

To facilitate practical use, we developed a user-friendly interactive online app (https://yunfeiwang.shinyapps.io/flu_characterization/) with the following functions: generating test datasets for testing, uploading datasets for testing, and providing a README. This app enables a simple and precise way of characterizing HA and NA types immediately after protein sequencing.

### HA subtype-predictive amino acids positions differ in sequence and structure

To understand the mechanism underlying the model’s predictive efficacy, we investigated the nature of the subtype-predictive reference residues and how their combined contribution drives subtype prediction. We analyzed one representative HA sequence from each subtype group (Supplementary Table 8), using H1 as the position-numbering reference for Group 1 and H3 for Group 2. Table 3 lists the amino acids used as references at positions 45, 48, 56, and 125 corresponding to the residues identified by the model as subtype-predictive.

**Table 3.**
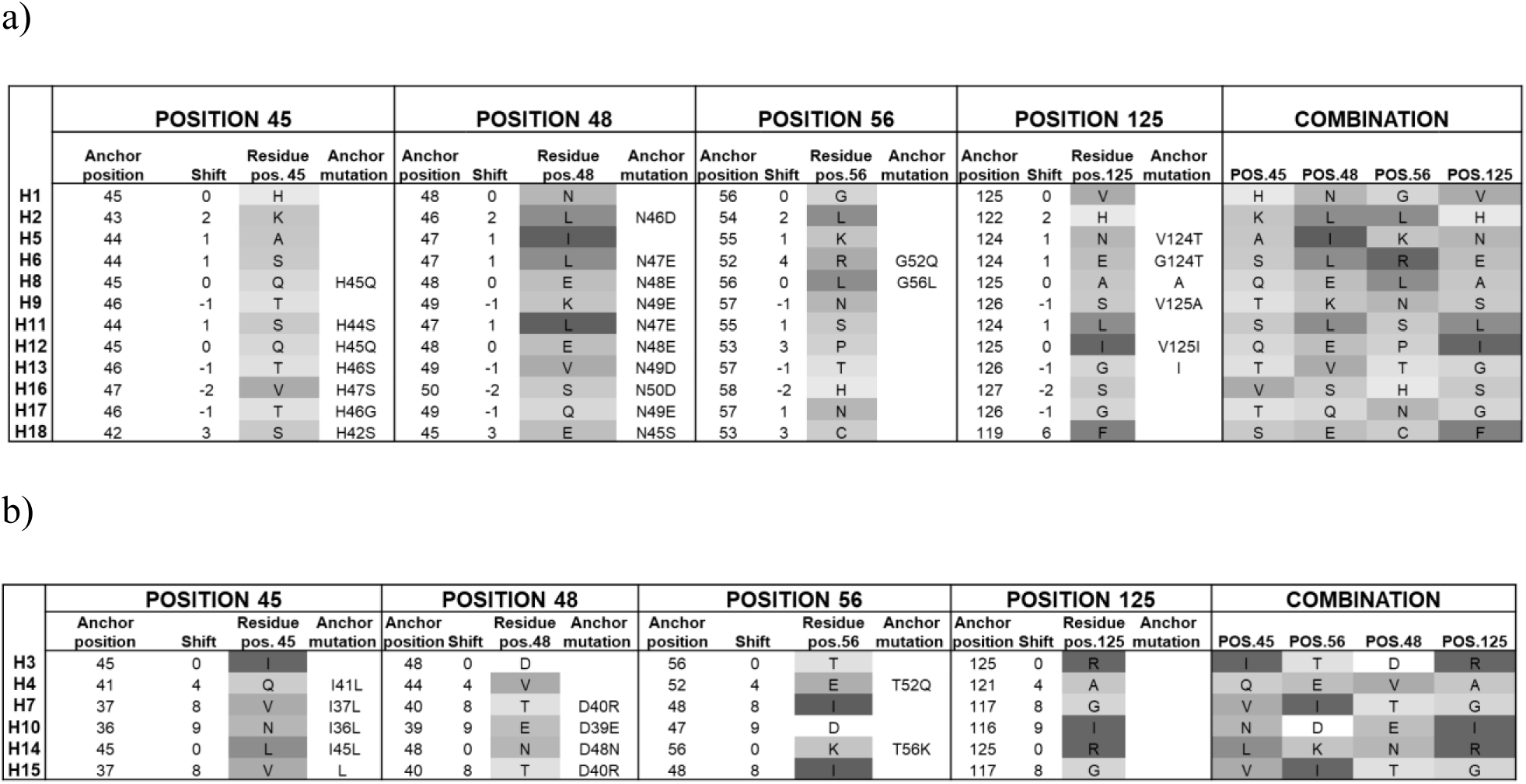
Position and shift of subtype-predictive residues and anchor residues for HA for group 1 (a) and group 2 (b).

Instead of conventional alignment, sequences were first superimposed onto H1 and H3 to preserve the amino acid indexing. The leader sequence peptide was also maintained in the initial evaluation. Table 3 shows that while H1 has histidine (pos. 45), asparagine (pos.48), glycine (pos. 56), and valine (pos. 125), the other subtypes have different amino acids at these positions. The combination of the four different amino acids is unique for each subtype, explaining how the model achieves subtype-prediction. The same phenomenon occurs for H3, which has isoleucine (pos. 45), aspartic acid (pos.48) threonine (pos. 56), and arginine (pos.125), while the rest of the subtypes in group 2 have different amino acids at these locations.

While being unique in H1 and H3 for these indexes numbered positions (shown for pos. 45 in Fig. 4A and 5A, respectively), these amino acids are generally conserved within the other HA subtypes but shifted in index numbering as analyzed above. This is confirmed structurally, as the subtype-predictive residues align in crystal structures of HAs (Fig.4D for group 1, Fig. 5D for group 2). The amino acid residues that appear in the same position in crystal structures we have identified as anchor residues, and these align structurally across different subtypes in each group despite having different index numbers. Thus, the combination of amino acid variation and position shifting results in the model being subtype-predictive. Positions that appear “variable” across subtypes represent a conserved anchor amino acid displaced by a subtype-specific offset. When the offset is the same across subtypes, the distinctiveness is conferred by a different amino acid.

**Fig. 4.**
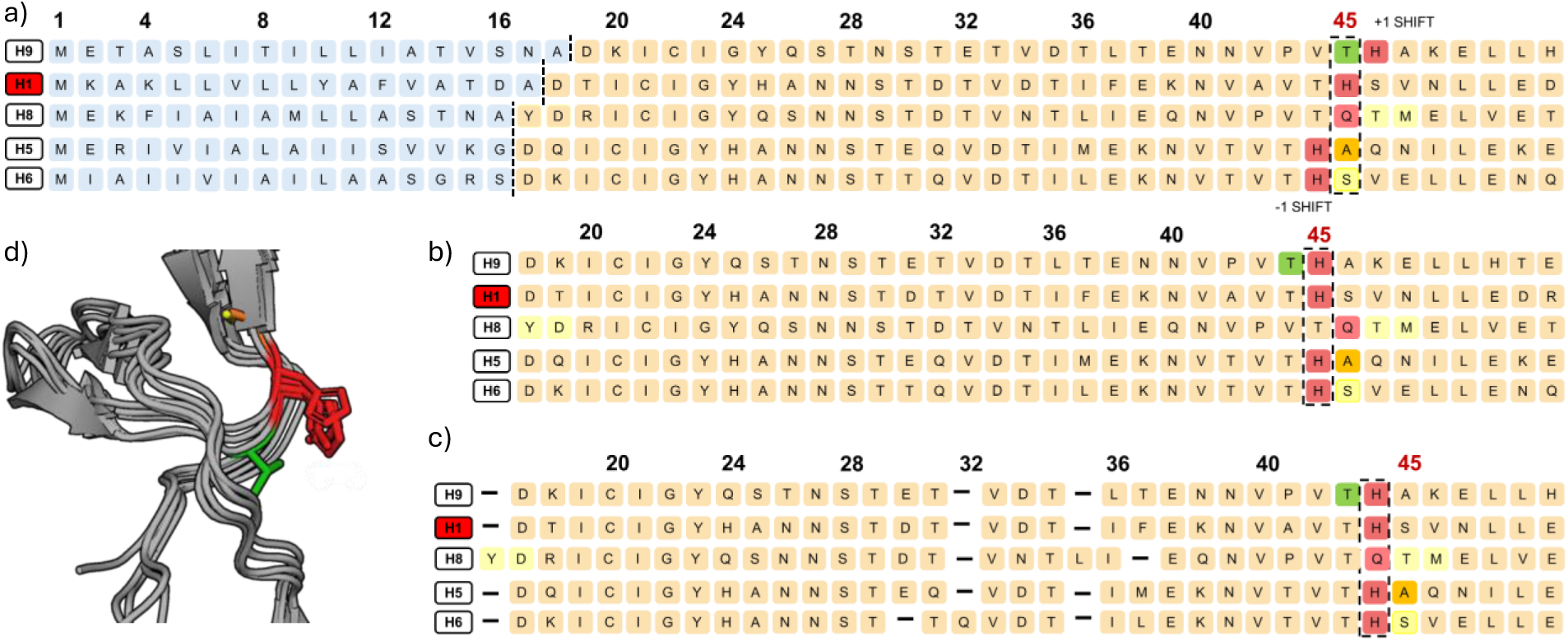
Sequence and structural location of VAR46 for group 1 HA.

**Fig. 5.**
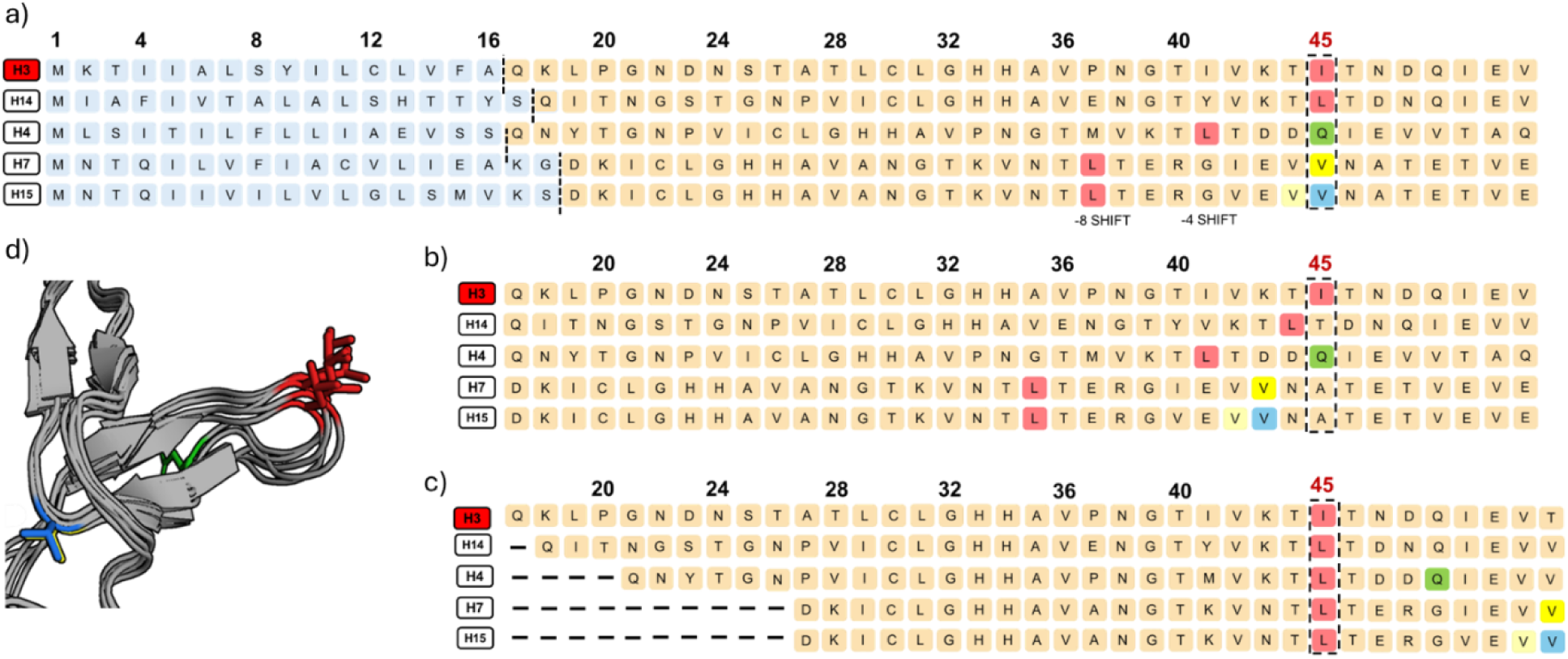
Sequence and structural location of VAR46 for group 2 HA.

### Leader Peptide length and Indels in HA contribute to the sequence shift of the anchor amino acids

To understand why anchor residues are displaced, we also examined signal peptide length for each sequence (Burke D.F. et al., 2014). Across all HA subtypes, the signal peptide is cleaved upon secretion, generating the mature HA sequence. While most HAs are cleaved between residues 16–17, substantial variation exists. Several subtypes exhibit cleavage at 14–15, 15–16, or 17–18, and signal peptide length can differ by 1–3 amino acids (Supplementary Table 8). In these cases, the N-terminal residue of the mature HA is shifted relative to the H1 reference. For group 1 HAs, this anchor residue is typically histidine, except for H8, where the aligned anchor is glutamine. When sequences are compared without signal peptides, all subtypes except H8 align at position 45. However, when indels are incorporated into the alignment, the apparent displacement disappears: all sequences, including H8, correctly align at reference position 45, restoring conservation of the anchor residue (Fig. 4 A, B, C). For group 2 HAs, this anchor residue is typically leucine, except for H3, where the aligned anchor is isoleucine. When sequences are compared without signal peptides, none of the subtypes aligns at position 45. However, when indels are incorporated into the alignment, the apparent displacement disappears: all sequences correctly align at the reference position 45, restoring conservation of the anchor residue (Fig. 5 A, B, C). Supplementary Fig.1 and 2 show the alignment of other anchor residues upon signal peptide removal and incorporation of indels for both group 1 and group 2. We located the subtype-predictive residues in crystal structures of HA from H1 and H3 (Fig. 6,7), showing how many predictive residues are localized in loops of HA. These positions represent the sequence variants whose predictive confidence is above 96% (Fig. 6,7 A), 94%(Fig. 6,7 B), 93%(Fig. 6,7 C). Supplementary Fig.3 and 4 show the distinctive localization of the other anchor-residues and the shift of the subtype-predictive residues.

**Fig. 6.**
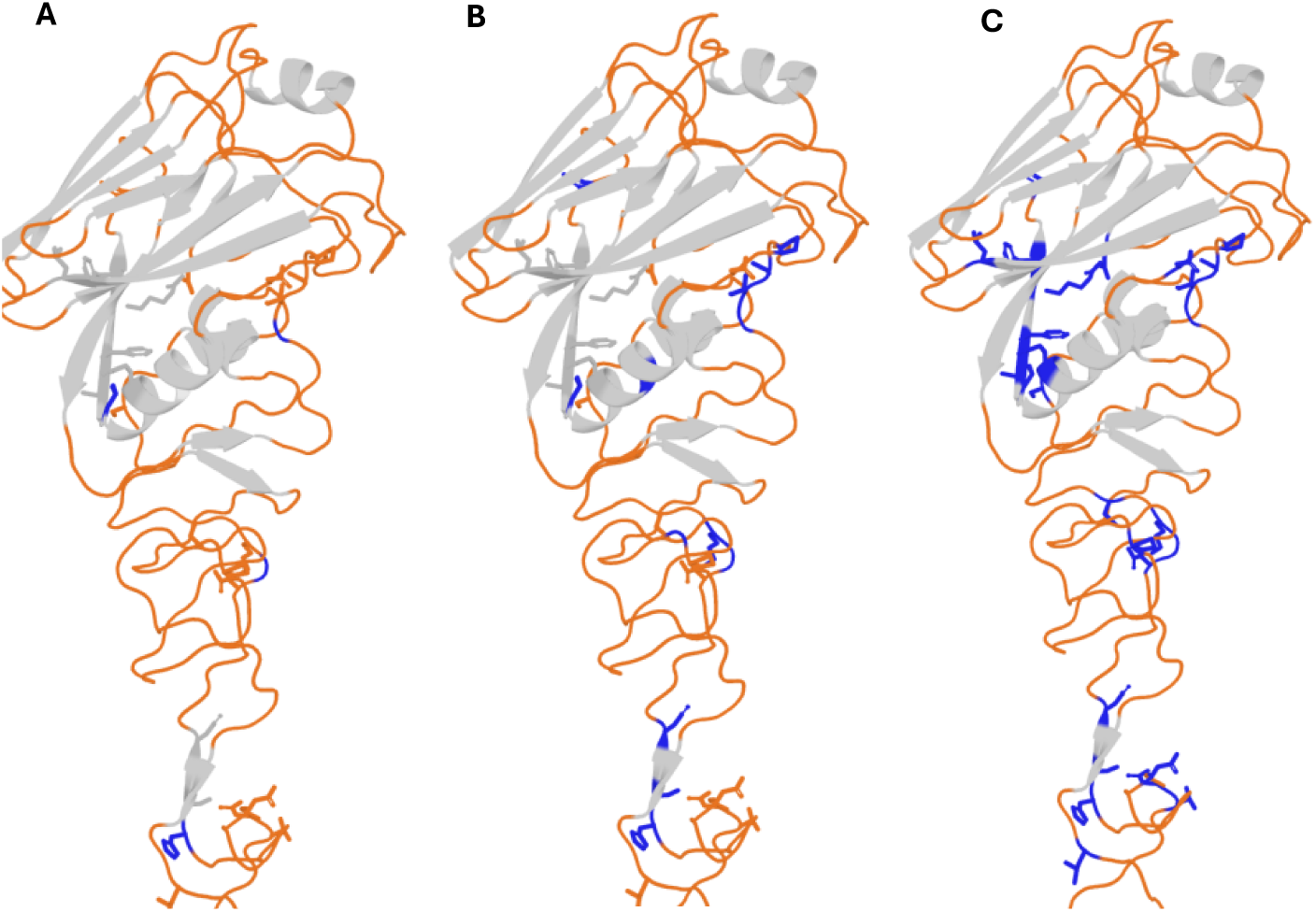
Structure localization of subtype-predictive amino acids in H1.

**Fig. 7.**
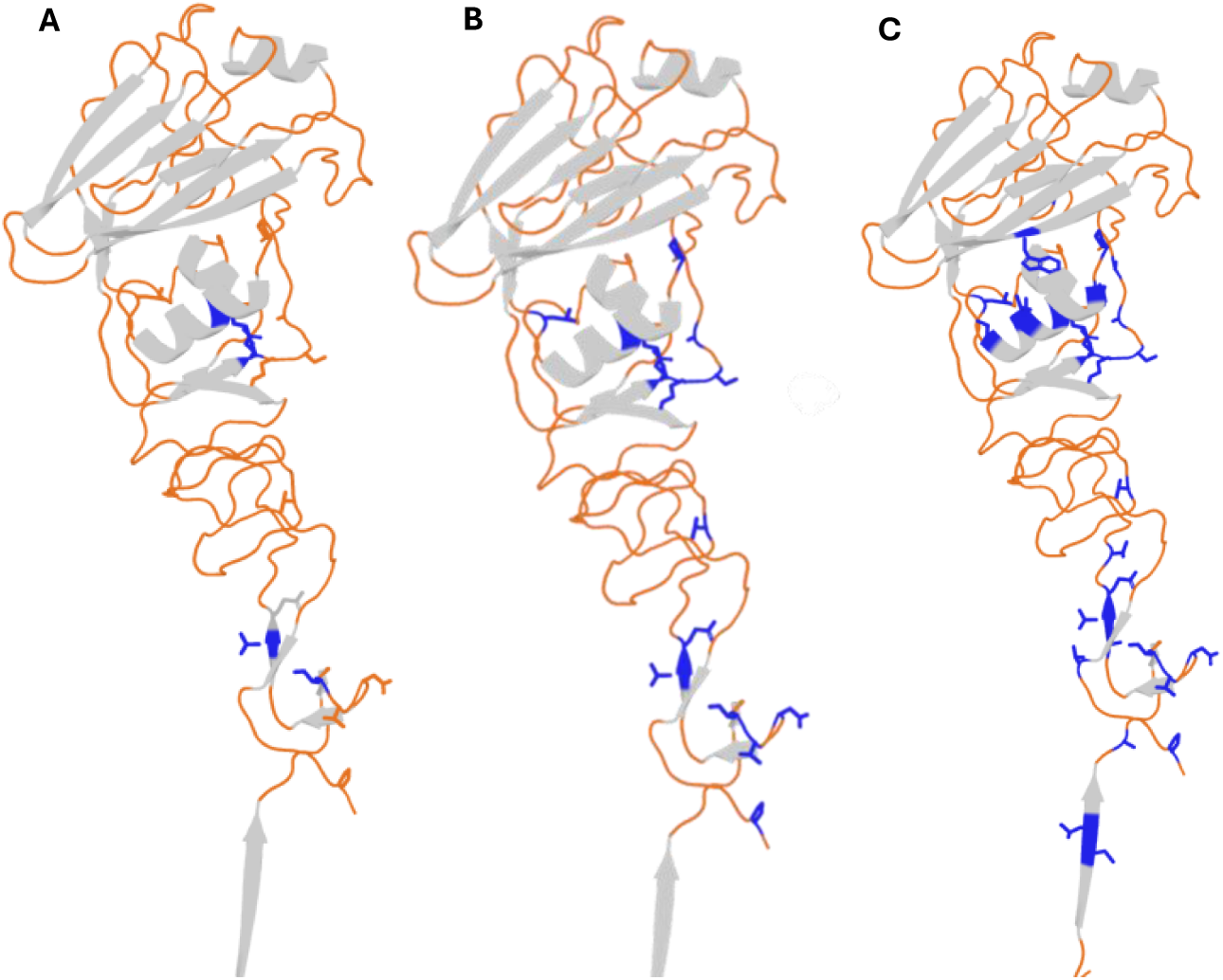
Structure localization of subtype-predictive amino acids in H3.

### Indels in NA contribute to the sequence shift of the anchor amino acids

We then determined whether the subtype-predictive reference residues worked for NA with the same mechanism we observed for HA sequences. We analyzed one representative NA sequence from each subtype group (Supplementary Table 9), using N1 as the position-numbering reference for Group 1 and N2 for Group 2. Table 4 lists the amino acids used as references at positions 34,45,111 and 409 corresponding to the residues identified by the model as subtype-predictive. Similarly to HA, instead of conventional alignment, NA sequences were first superimposed onto N1 and N2 to preserve the amino acid indexing. However, unlike HA, anchor amino acids are less conserved within the other NA subtypes. Once the anchors were identified structurally (Fig.8, 9 C), we determined the presence of a shift in index numbering. However, given the absence of a signal peptide in NA, the shift is solely attributable to indels (Fig.8, 9 A and B). Supplementary Fig.5 and 6 show the alignment of other anchor residues upon incorporation of indels for both group 1 and group 2. Supplementary Fig.7 and 8 shows the localization of anchor 409 and the respective shift of subtype predictive residues on the structure.

**Fig. 8.**
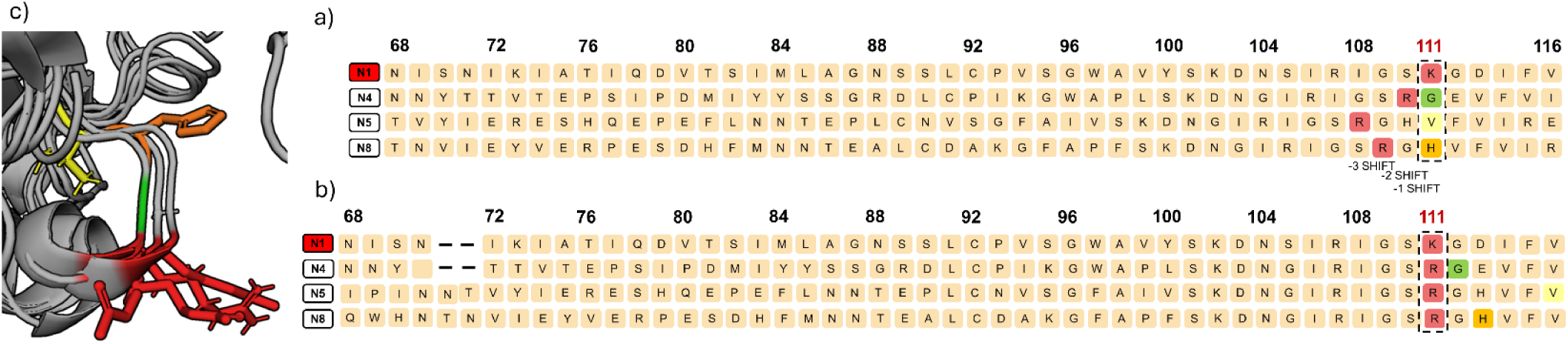
Sequence and structural location of Position 111 for group 1 NA.

**Fig. 9.**
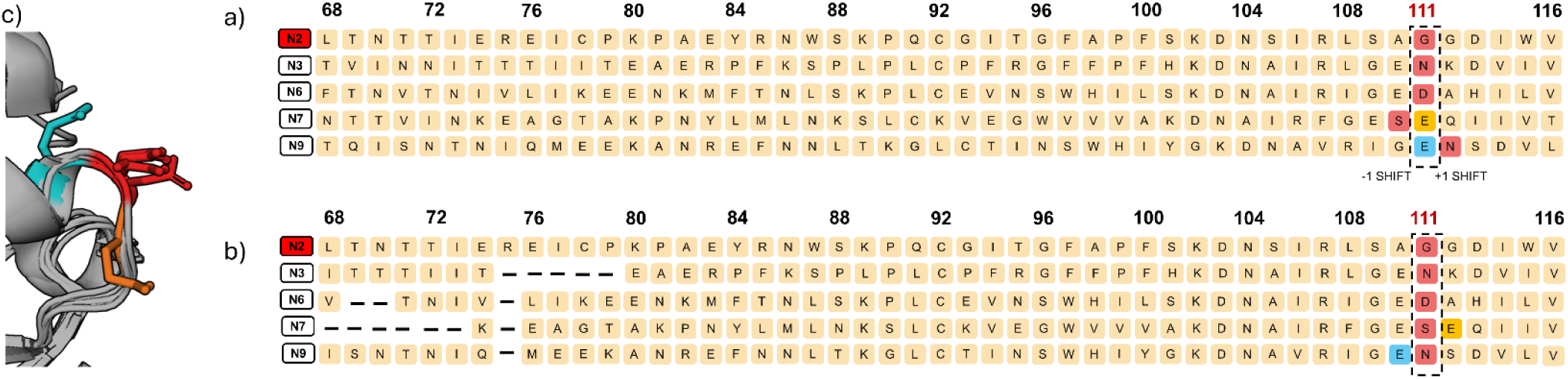
Sequence and structural location of Position 111 for group 2 NA.

**Table 4.** Position and shift of subtype-predictive residues and anchor residues for NA.

|  | POSITION 34 |  |  |  | POSITION 45 |  |  |  | POSITION 111 |  |  |  | POSITION 409 |  |  |  | COMBINATION |  |  |  |
| --- | --- | --- | --- | --- | --- | --- | --- | --- | --- | --- | --- | --- | --- | --- | --- | --- | --- | --- | --- | --- |
|  | Anchor position | Shift | Residue pos. 34 | Anchor mutation | Anchor position | Shift | Residue pos. 45 | Anchor mutation | Anchor position | Shift | Residue pos. 111 | Anchor mutation | Anchor position | Shift | Residue pos. 409 | Anchor mutation | POS.34 | POS.45 | POS.111 | POS.409 |
| N1 | 34 | 0 | I |  | 45 | 0 | H |  | 111 |  | K |  | 409 |  | H |  | I | H | K | H |
| N4 | 34 | 0 | F | I34F | 44 | 1 | E |  | 110 | 1 | G | K110R | 410 | -1 | I | R | F | E | G | I |
| N5 | 34 | 0 | I |  | 42 | 3 | T | H42V | 108 | 3 | V | K108R | 411 | -2 | T | P | I | T | V | T |
| N8 | 34 | 0 | L | I34L | 41 | 4 | C | C41N | 109 | 2 | H | K109R | 409 | 0 | P | P | L | C | H | P |

b)
|  | POSITION 34 |  |  |  | POSITION 45 |  |  |  | POSITION 111 |  |  |  | POSITION 409 |  |  |  | COMBINATION |  |  |  |
| --- | --- | --- | --- | --- | --- | --- | --- | --- | --- | --- | --- | --- | --- | --- | --- | --- | --- | --- | --- | --- |
|  | Anchor position | Shift | Residue pos. 34 | Anchor mutation | Anchor position | Shift | Residue pos. 45 | Anchor mutation | Anchor position | Shift | Residue pos. 111 | Anchor mutation | Anchor position | Shift | Residue pos. 409 | Anchor mutation | POS.34 | POS.45 | POS.111 | POS.409 |
| N2 | 34 | 0 | T |  | 45 | 0 | P |  | 111 |  | G |  | 409 | 0 | I |  | T | P | G | I |
| N3 | 34 | 0 | V |  | 45 | 0 | V |  | 111 | 0 | N | G111N | 408 | 1 | F | I408S | V | V | N | F |
| N6 | 34 | 0 | G |  | 45 | 0 | T |  | 111 | 0 | D | G111D | 409 | 0 | A | I409A | G | T | D | A |
| N7 | 34 | 0 | S |  | 45 | 0 | D |  | 110 | 1 | E | G110S | 408 | 1 | F | I408S | S | D | E | F |
| N9 | 34 | 0 | G |  | 45 | 0 | N |  | 112 | -1 | E | G112N | 409 | 0 | S | I409S | G | N | E | S |

We located the subtype-predictive residues in crystal structures of NA from N1 and N2 (Fig. 10,11), showing how two of the 4 predictive residues are localized in loops of the protein. We could not localize the location of position 34 and 45 as the N-terminus portion of the protein is not included in the crystallized format.

**Fig. 10.**
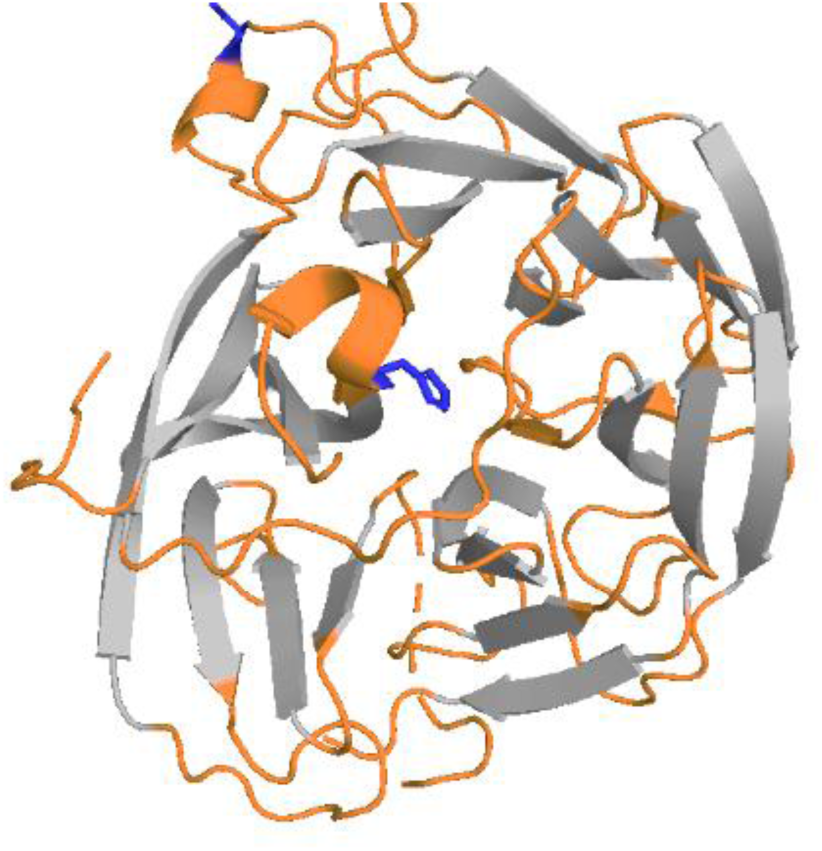
Structure localization of subtype-predictive amino acids in N1.

**Fig. 11.**
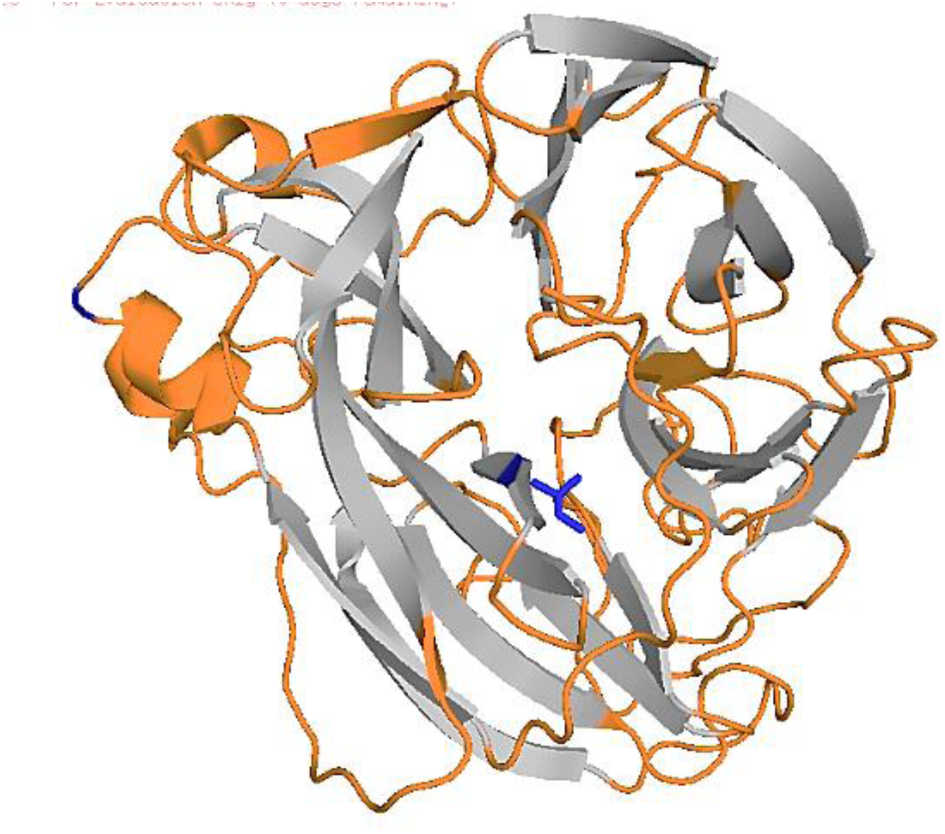
Structure localization of subtype-predictive amino acids in N2.

Supplementary Fig.7 and 8 show the distinctive localization of the other anchor-residues and the shift of the subtype-predictive residues.

## Discussion

Several machine learning methods have been developed for influenza A virus subtype classification. Humayun et al. (Humayun et al., 2021), for instance, achieved high accuracy using avian flu nucleotide sequence information and physicochemical properties with an accuracy of 0.95 for HA. More recently, researchers have explored a variety of approaches. WaveSeekerNet uses an attention-based deep learning model to predict both HA/NA subtype (Nguyen, H.H., et al., 2025.). He and colleagues (He, L., et al. 2024) developed SCCFV, an alignment-free method that combined with random forest reportedly hits 99.98% accuracy for HA and NA subtyping. PreIS, meanwhile, takes a protein language model approach but is limited to HA-only prediction with an accuracy of 0.95 (Sohrabi, M.A., et al. 2024).

These existing methods have a few limitations. Some are trained primarily on avian sequences, so it is unclear how well they generalize to human or other mammalian hosts. Also, some focus on HA subtyping alone—PreIS is a good example—while NA subtyping remains underexplored. Beyond that, many of these approaches rely on complex feature engineering or black-box deep learning models, which can make them difficult to interpret biologically. WaveSeekerNet and Flu-CNN fall into this category.

Our method takes a different approach. We use protein sequences rather than nucleotides, and include viruses from diverse hosts—humans, birds, swine, and others. Our model handles both HA and NA subtypes within the same framework and does so with near-perfect accuracy. And because we built it using a straightforward logistic regression approach, we can trace predictions back to specific amino acid positions. This makes the model not just accurate but specific and meaningful, which leads to our further structural study in this paper.

Additionally, we introduce a user-friendly interactive online application for a simple and precise way of characterizing HA and NA subtypes directly following protein sequence acquisition. This application successfully predicted the circulating H5N1 strain responsible for conjunctivitis in Texan ranch hands in 2024 and predicted two independent large HA and NA datasets with almost perfect accuracies (0.9989 for HA, and 0.9987 for NA). Overall, the advancement of sequence techniques combined with the technique reported here makes it easy, simple, precise, and fast to identify HA and NA subtypes.

Our analysis identified subtype-predictive amino acids for HA and NA subtypes using a large dataset, allowing us to have strong statistical estimations and data mining for model building. However, metadata inaccuracies on source organisms and dates might exist. We grouped subtypes H8, H12, and H14-H18 due to their low frequency, thus we will need to revisit these subtypes as additional data become available. Similarly, N10 and N11 were excluded from the model fitting due to the limited number of available cases. As new sequence information becomes available from global surveillance, it is possible that new subtypes will be identified and that the model will need to be refined. We note that this problem is the same for other methods where antigens and antisera do not yet exist.

Our model was able to exploit a difference in signal peptide leader length and indels for HA and indels for NA to identify a shift of conserved amino acids in specific locations. This allowed us to identify at specific subtype-predictive locations a set of amino acids that were unique for each subtype. By analyzing HA and NA sequences using alternative representations rather than relying exclusively on conventional sequence alignment, we detected informative patterns that were otherwise obscured in conventional analysis. This strategy allows the identification of hidden sequence signals, such as subtype-specific fingerprints, which are not obvious when residues are aligned.

In conclusion, these findings pave a simple way for improved HA/NA subtype classification, potentially contributing to enhanced influenza surveillance and control measures.

## Materials and Methods

### Datasets

Two datasets were used, comprising HA and NA protein sequence data. The protein sequences of the HA and NA were downloaded from GISAID EpiFlu (Shu Y. et al., 2017) on June 5^th^, 2019. Each dataset contains two variables: a full-length NA or HA protein sequence and an ID, from which the strain, HA type, NA type, host, and year can be extracted. The database includes 52.6% human strains and 47.4% animal strains spanning from 1902 to 2019. Each HA protein sequence comprises 577 amino acids, while each NA sequence contains 477 amino acids.

### Statistical Analysis

Statistical analysis was conducted using R version 4.2.3, utilizing the “nnet” and “randomForest” packages. Initially, a single variable predictor logistic regression model was fitted to identify the top variables to fit the final HA or NA prediction models. Subsequently, multiple-variable logistic regression and random forest models were fitted, and an online R Shiny app was developed.

### Logistic Regression Model

The logistic regression prediction model is simple, particularly in relation to the odds ratio. The probability for sequence *Y_i_* belonging to subtype *k* from the multinomial logistic regression model is defined as follows:

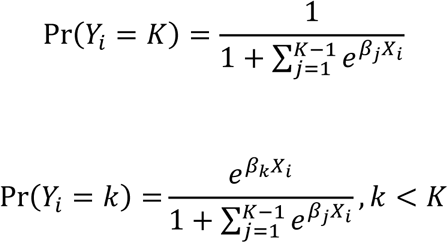

where *Y_i_* represents the HA or NA subtype for subject *i*, *β_j_* denotes the coefficient associated with the subtype *j*, and *X^i^* is the predictor.

### Random Forest

Random forest is an ensemble learning algorithm for classification composed of multiple decision trees, with each tree voting on the most popular class. The algorithm involves three key steps. First, it employs random feature selection, randomly selecting features at each split. Second, it uses bootstrap aggregating (bagging), building trees on bootstrapped subsets. Third, it applies an ensemble method, aggregating predictions from multiple trees to produce the final classification.

### Evaluation

Model evaluation was based on accuracy (ACC), precision (PREC), recall (REC), and F1 score. The F1 score is calculated as the harmonic mean of precision and recall. The formulas are as follows:

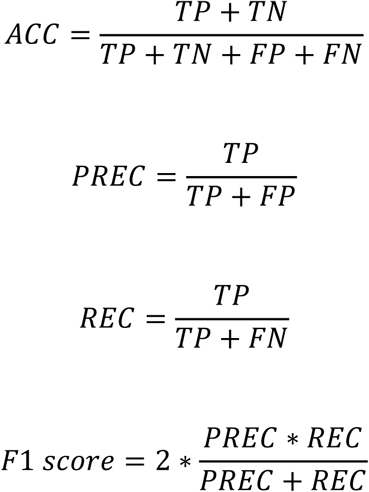

where TP, TN, FP, and FN stand for true positive, true negative, false positive, and false negative.

### Cross-validation

An 80/20 train–test split was used for cross validation. The data were randomly split into ten parts, with eight parts used as the training dataset and the remaining two parts as the test dataset. Each time, a logistic regression model or a random forest model was constructed using the training dataset, and predictions were subsequently made on the test dataset. The metrics scores, such as accuracy (ACC), precision (PREC), recall (REC), and F1 score based on the predictions on the test dataset, were recorded. This process was repeated 1000 times, and the average scores from these repetitions were considered the final scores.

### Independent validation

We used two additional independent HA and NA protein datasets collected from 2020 to 2024. The HA protein dataset contains 23,231 observations and includes 11 major HA subtypes and 7 minor subtypse (H8, H12, H14-18). We use the umbrella, designation “H99” to signify any of the seven minor HA subtypes. The NA protein dataset contains 23,133 observations and includes 9 NA subtypes (N1-N9) from 2020 to 2024. Simply uploading the HA or NA protein fasta file to the app at https://yunfeiwang.shinyapps.io/flu_characterization/, and by clicking the run characterization button, we can see the predicted subtype HA and NA on the output screen, or can download the results by clicking the download button. The accuracy of the predictions was estimated as the percentage of the match of the predicted subtype and the actual subtype.

### Sequence and structure analysis

Sequence alignment was performed using Clustal Omega (Sievers F. et al., 2018). Identification of signal peptide cleavage site was done using SignalP 6.0 (Nielsen H. et al., 2025). Structure analysis was performed using Pymol. PDBs for HA used were the following from group 1: subtype 1 (PDB:1rvz), subtype 5 (PDB: 3zp0), subtype 6 (PBD:5T0B), subtype 8 (PDB:6ZRK), subtype 9 (PDB: 1JDS); and group 2: subtype 3 (PDB:9BDF), subtype 4 (PDB:5Y2M), subtype 7 (PBD:6II9), subtype 14 (PDB:6V48), subtype 15 (PDB:5TG9). PDBs for NA used were the following from group 1: subtype 1 (PDB:7U2Q), subtype 4 (PDB:2HTV), subtype 5 (PBD:9EIT), subtype 8 (Alphafold3 prediction of sequence A/Mallard/Utah/D1615280/2016/ H7N8 using online portal https://alphafoldserver.com); and group 2: subtype 2 (PDB:9MQW), subtype 3 (PDB:4HZV), subtype 6 (PBD:6HGB), subtype 7 (PDB:4QN3), subtype 9 (PDB:6HFC).

## Availability and Implementation

An online application is available at https://yunfeiwang.shinyapps.io/flu_characterization/

## Supplementary information

Supplementary data are available at *Bioinformatics* online.

## Acknowledgements

The authors would like to thank Kevin R. McCarthy for his invaluable support and guidance throughout this work.

## Authors contributions

Y.W. performed the statistical analysis, predictive modeling, and validation of results. F.V. performed the validation with sequences and structures. Two additional independent HA and NA protein datasets collected from 2020 to 2024 were provided by M.M.H and K.W.

Y.W, F.V. and M.A.M. wrote the draft of the manuscript. Y.W and F.V. developed the figures. M.A.M. secured the funding. M.A.M. and R.W.R. supervised the work. All authors reviewed and approved the final version of the manuscript.

## Data and Code availability statement

The codes for flu subtype characterization were implemented in R 4.3.0. Data and codes are publicly available at Github: https://github.com/wang-484/flu_characterization.

---

**Supplementary Fig. 1.**
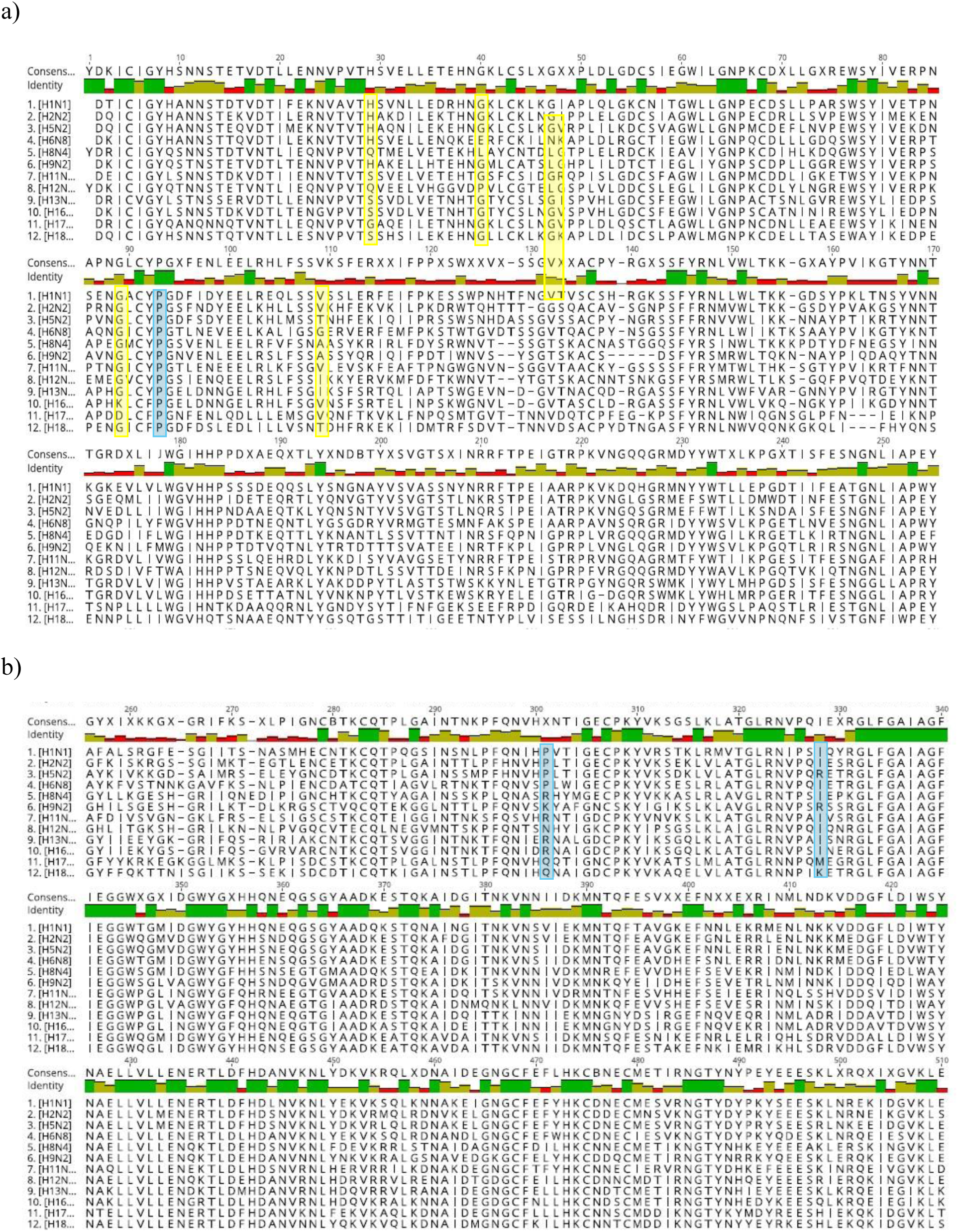
Crystal structure of HA stem belonging to subtype 1 (PDB:1rvz), subtype 5 (PDB: 3zp0), subtype 6 (PBD:5T0B), subtype 8 (PDB:6ZRK), subtype 9 (PDB: 1JDS). The conserved anchor residue is highlighted in red, while the neighboring reference residues are colored in orange (subtype 5), yellow (subtype 6), green (subtype 9).

**Supplementary Fig. 2.**
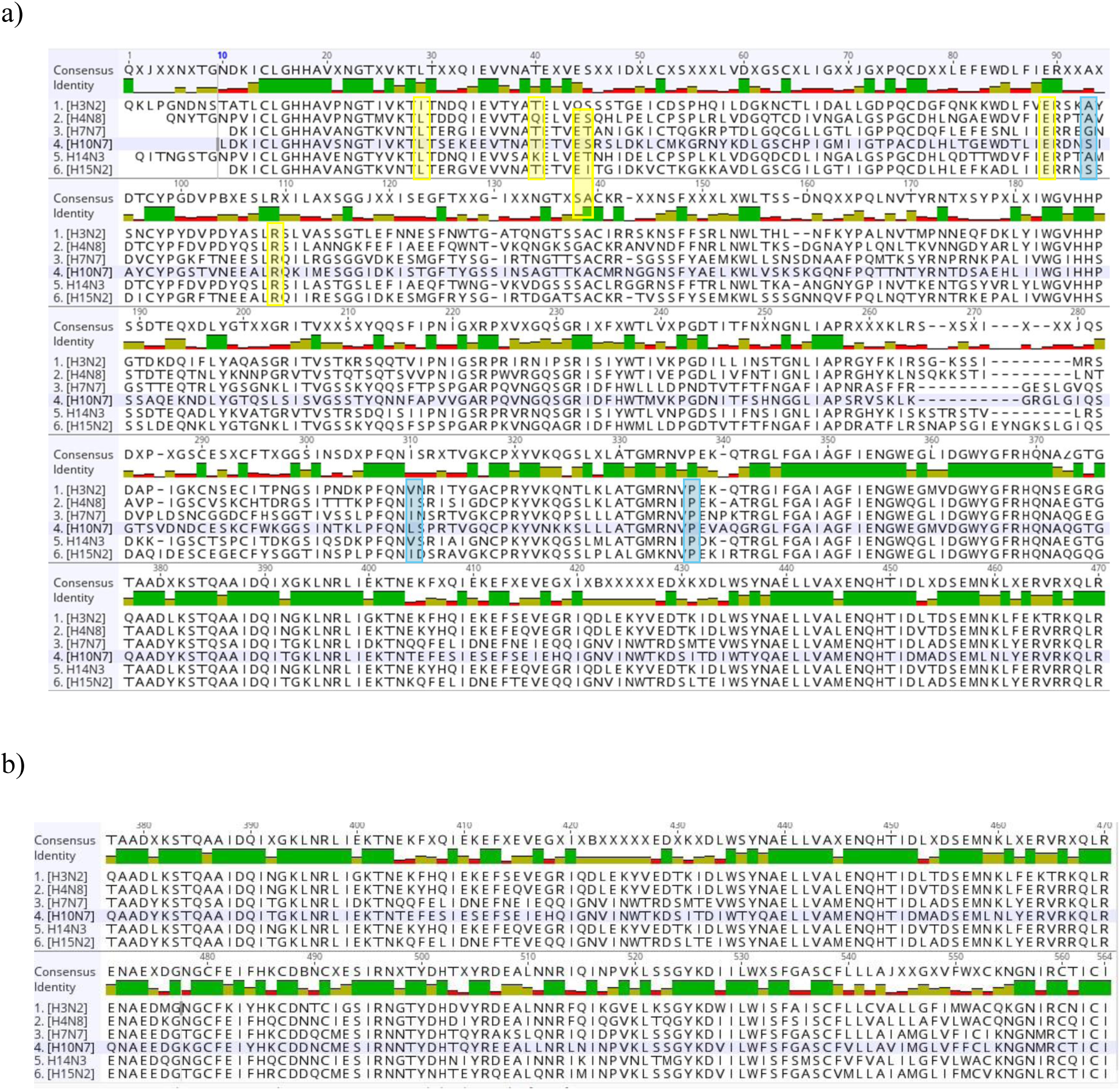
Highlight of anchor residues aligned in the subtype-predictive location upon removal of signal peptide and indels incorporation for group 1 HA sequences. Yellow highlights subtype-predictive residues locations of confidence level of 96%, while blue 94%.

**Supplementary Fig. 3.**
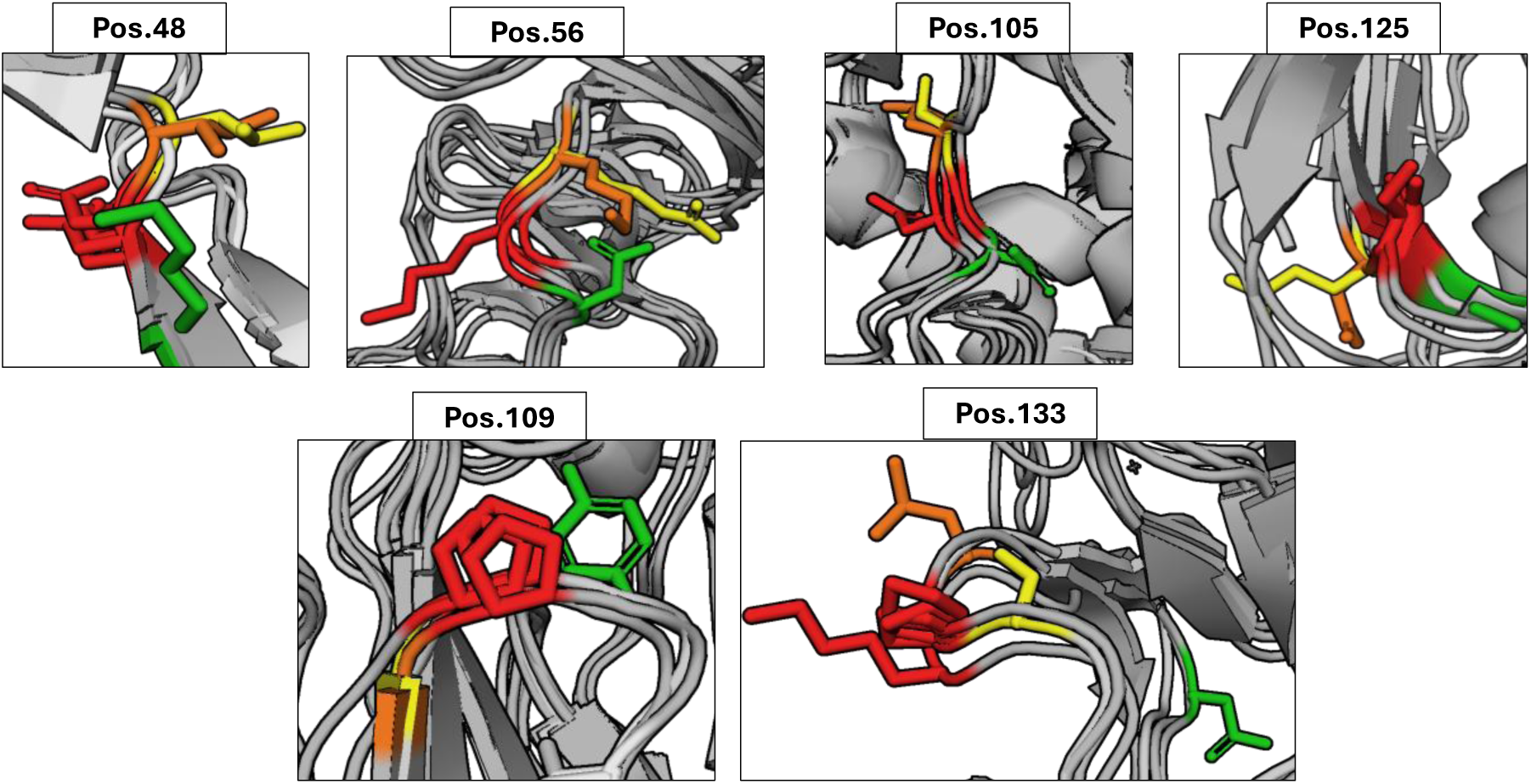
Highlight of anchor residues aligned in the subtype-predictive location upon removal of signal peptide and indels incorporation for group 2 HA sequences. Yellow highlights subtype-predictive residues locations of confidence level of 96%, while blue 94%.

**Supplementary Fig. 4.**
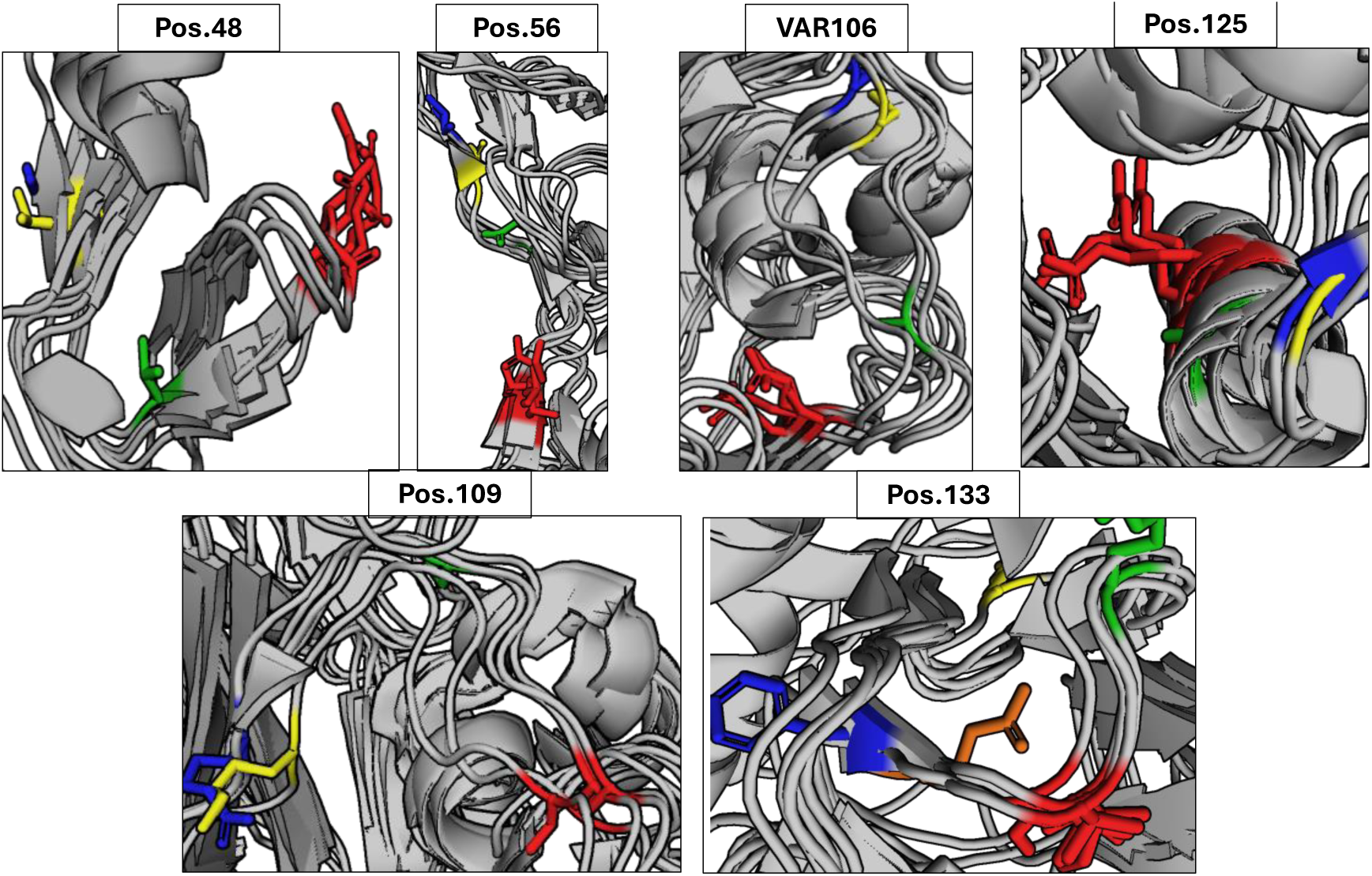
Crystal structure of HA stem belonging to subtype 3 (PDB:9BDF), subtype 4 (PDB:5Y2M), subtype 7 (PBD:6II9), subtype 14 (PDB:6V48), subtype 15 (PDB:5TG9). The conserved anchor residue is highlighted in red, while the neighboring reference residues are colored in orange (subtype 14), blue (subtype 15), yellow (subtype 7), green (subtype 4).

**Supplementary Fig. 5.**
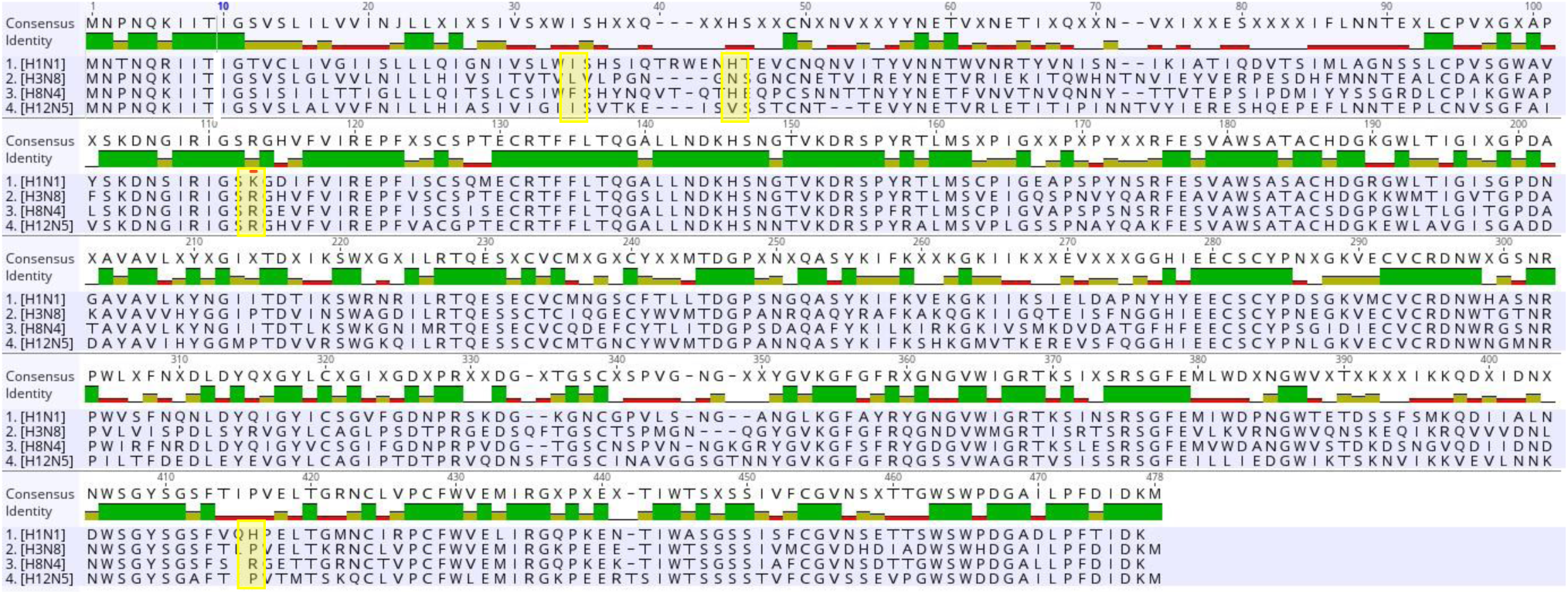
Highlight of anchor residues aligned in the subtype-predictive location upon removal of signal peptide and indels incorporation for group 1 NA sequences. Yellow highlights subtype-predictive residues locations of confidence level of 96%.

**Supplementary Fig. 6.**
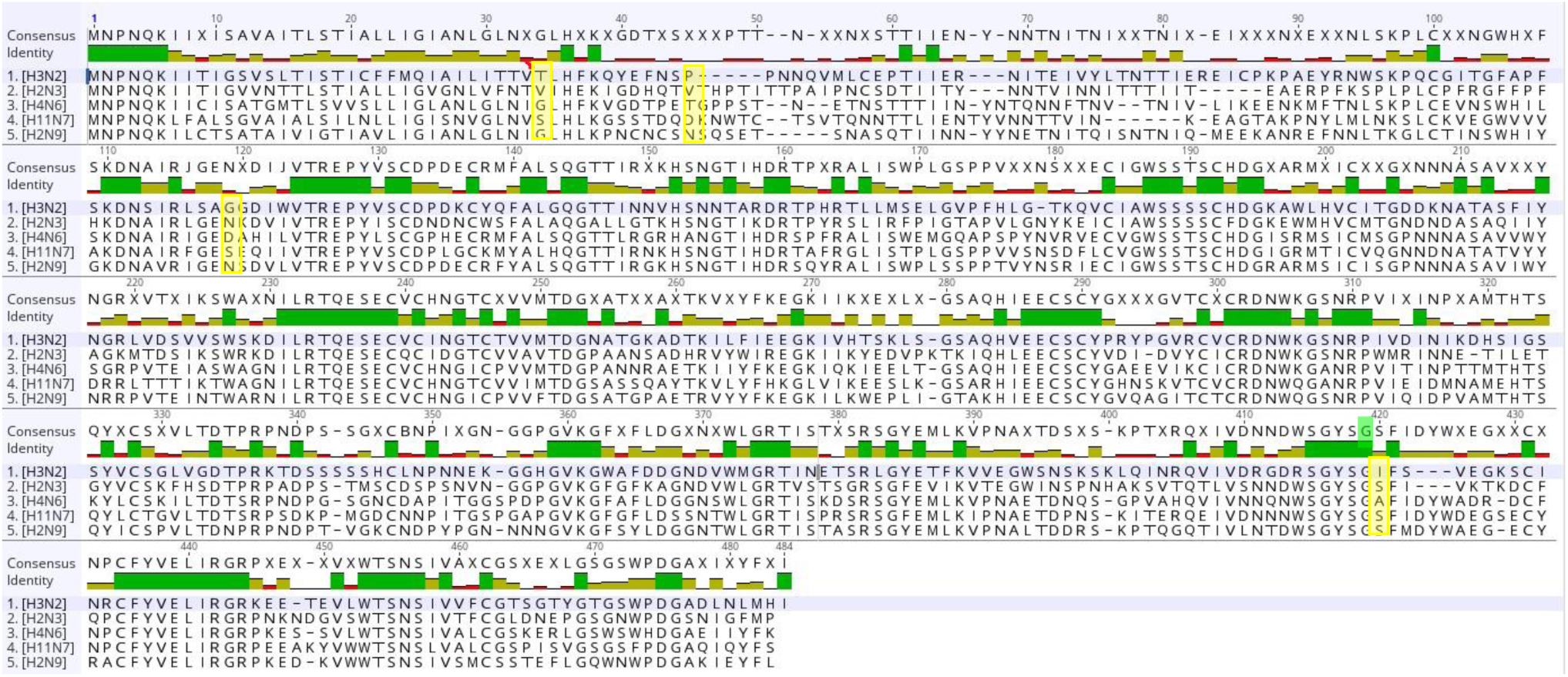
Highlight of anchor residues aligned in the subtype-predictive location upon removal of signal peptide and indels incorporation for group 2 NA sequences. Yellow highlights subtype-predictive residues locations of confidence level of 96%.

**Supplementary Fig. 7.**
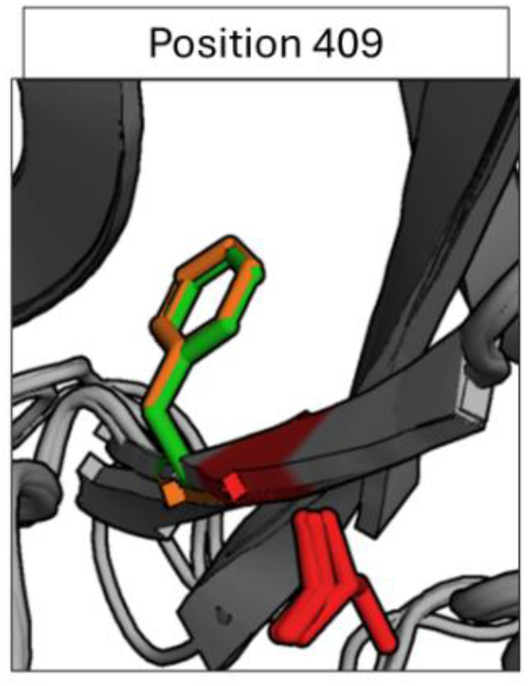
Crystal structure of NA belonging to subtype 2 (PDB:9MQW), subtype 3 (PDB:4HZV), subtype 6 (PBD:6HGB), subtype 7 (PDB:4QN3), subtype 9 (PDB:6HFC). The conserved anchor residue is highlighted in red, while the neighboring reference residues are colored in green (subtype 3), orange (subtype 7).

**Supplementary Fig. 8.**
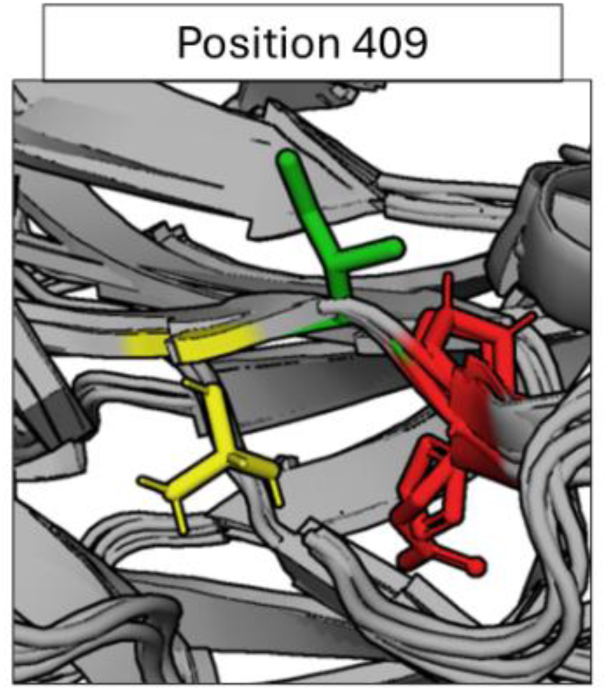
Crystal structure of NA belonging to subtype 1 (PDB:7U2Q), subtype 4 (PDB:2HTV), subtype 5 (PBD:9EIT), subtype 8 (Alphafold prediction of…). The conserved anchor residue is highlighted in red, while the neighboring reference residues are colored in green (subtype 4), yellow (subtype 5).

**Supplementary Table 1s.** The distribution of amino acids in HA Position 56 among different HA types.

| strain | Pos56 |  |  |  |  |  |  |  |  |  |  |  |  |  |  |  |  |  |  |  |  |
| --- | --- | --- | --- | --- | --- | --- | --- | --- | --- | --- | --- | --- | --- | --- | --- | --- | --- | --- | --- | --- | --- |
|  | A | C | D | E | F | G | H | I | K | L | M | N | P | R | S | T | V | W | X | Y | Total |
| 1 | 0 | 0 | 2 | 69 | 0 | 30277 | 0 | 0 | 1 | 1 | 0 | 67 | 0 | 35 | 1 | 0 | 7 | 1 | 4 | 1 | 30466 |
|  | 0 | 0 | 0.01 | 0.23 | 0 | 99.38 | 0 | 0 | 0 | 0 | 0 | 0.22 | 0 | 0.11 | 0 | 0 | 0.02 | 0 | 0.01 | 0 |  |
| 10 | 0 | 0 | 916 | 0 | 0 | 14 | 0 | 0 | 1 | 0 | 0 | 363 | 0 | 0 | 0 | 0 | 0 | 0 | 0 | 0 | 1294 |
|  | 0 | 0 | 70.79 | 0 | 0 | 1.08 | 0 | 0 | 0.08 | 0 | 0 | 28.05 | 0 | 0 | 0 | 0 | 0 | 0 | 0 | 0 |  |
| 11 | 0 | 0 | 0 | 0 | 0 | 0 | 0 | 0 | 0 | 0 | 0 | 0 | 0 | 0 | 715 | 1 | 0 | 0 | 0 | 0 | 716 |
|  | 0 | 0 | 0 | 0 | 0 | 0 | 0 | 0 | 0 | 0 | 0 | 0 | 0 | 0 | 99.86 | 0.14 | 0 | 0 | 0 | 0 |  |
| 13 | 0 | 0 | 0 | 0 | 0 | 0 | 0 | 0 | 0 | 0 | 0 | 0 | 0 | 0 | 0 | 582 | 0 | 0 | 0 | 0 | 582 |
|  | 0 | 0 | 0 | 0 | 0 | 0 | 0 | 0 | 0 | 0 | 0 | 0 | 0 | 0 | 0 | 100 | 0 | 0 | 0 | 0 |  |
| 2 | 0 | 0 | 0 | 4 | 0 | 0 | 0 | 0 | 0 | 626 | 0 | 0 | 0 | 0 | 0 | 0 | 0 | 0 | 0 | 0 | 630 |
|  | 0 | 0 | 0 | 0.63 | 0 | 0 | 0 | 0 | 0 | 99.37 | 0 | 0 | 0 | 0 | 0 | 0 | 0 | 0 | 0 | 0 |  |
| 3 | 60 | 0 | 0 | 228 | 0 | 0 | 0 | 8 | 0 | 0 | 0 | 2 | 0 | 0 | 0 | 29298 | 0 | 0 | 0 | 0 | 29596 |
|  | 0.2 | 0 | 0 | 0.77 | 0 | 0 | 0 | 0.03 | 0 | 0 | 0 | 0.01 | 0 | 0 | 0 | 98.99 | 0 | 0 | 0 | 0 |  |
| 4 | 0 | 0 | 0 | 1943 | 0 | 0 | 0 | 0 | 0 | 0 | 0 | 0 | 0 | 0 | 1 | 0 | 3 | 0 | 0 | 0 | 1947 |
|  | 0 | 0 | 0 | 99.79 | 0 | 0 | 0 | 0 | 0 | 0 | 0 | 0 | 0 | 0 | 0.05 | 0 | 0.15 | 0 | 0 | 0 |  |
| 5 | 0 | 0 | 0 | 0 | 0 | 0 | 2 | 0 | 5539 | 0 | 5 | 6 | 0 | 448 | 0 | 2 | 0 | 0 | 0 | 0 | 6002 |
|  | 0 | 0 | 0 | 0 | 0 | 0 | 0.03 | 0 | 92.29 | 0 | 0.08 | 0.1 | 0 | 7.46 | 0 | 0.03 | 0 | 0 | 0 | 0 |  |
| 6 | 0 | 0 | 0 | 0 | 0 | 0 | 0 | 2 | 4 | 0 | 0 | 0 | 0 | 1811 | 0 | 0 | 0 | 0 | 0 | 0 | 1817 |
|  | 0 | 0 | 0 | 0 | 0 | 0 | 0 | 0.11 | 0.22 | 0 | 0 | 0 | 0 | 99.67 | 0 | 0 | 0 | 0 | 0 | 0 |  |
| 7 | 0 | 0 | 0 | 2 | 6 | 0 | 0 | 2151 | 0 | 6 | 3 | 0 | 0 | 1 | 3 | 121 | 515 | 0 | 0 | 0 | 2808 |
|  | 0 | 0 | 0 | 0.07 | 0.21 | 0 | 0 | 76.6 | 0 | 0.21 | 0.11 | 0 | 0 | 0.04 | 0.11 | 4.31 | 18.34 | 0 | 0 | 0 |  |
| 9 | 0 | 12 | 1 | 0 | 0 | 0 | 7 | 1 | 1 | 0 | 3 | 7164 | 0 | 0 | 0 | 2 | 0 | 0 | 0 | 0 | 7191 |
|  | 0 | 0.17 | 0.01 | 0 | 0 | 0 | 0.1 | 0.01 | 0.01 | 0 | 0.04 | 99.62 | 0 | 0 | 0 | 0.03 | 0 | 0 | 0 | 0 |  |
| 99 | 1 | 2 | 0 | 0 | 0 | 0 | 269 | 21 | 37 | 1 | 0 | 3 | 478 | 1 | 6 | 4 | 0 | 0 | 0 | 0 | 823 |
|  | 0.12 | 0.24 | 0 | 0 | 0 | 0 | 32.69 | 2.55 | 4.5 | 0.12 | 0 | 0.36 | 58.08 | 0.12 | 0.73 | 0.49 | 0 | 0 | 0 | 0 |  |
| Total | 61 | 14 | 919 | 2246 | 6 | 30291 | 278 | 2183 | 5583 | 634 | 11 | 7605 | 478 | 2296 | 726 | 30010 | 525 | 1 | 4 | 1 | 83872 |
|  | 0.07 | 0.02 | 1.1 | 2.68 | 0.01 | 36.12 | 0.33 | 2.6 | 6.66 | 0.76 | 0.01 | 9.07 | 0.57 | 2.74 | 0.87 | 35.78 | 0.63 | 0 | 0 | 0 | 100 |

**Supplementary Table 2s.** The distribution of amino acids in HA Position 45 among different HA types.

| strain | Pos45 |  |  |  |  |  |  |  |  |  |  |  |  |  |  |  |  |  |  |
| --- | --- | --- | --- | --- | --- | --- | --- | --- | --- | --- | --- | --- | --- | --- | --- | --- | --- | --- | --- |
|  | A | D | E | F | G | H | I | K | L | N | P | Q | R | S | T | V | X | Y | Total |
| **1** | 0 | 0 | 0 | 0 | 0 | 30376 | 1 | 0 | 0 | 2 | 2 | 2 | 3 | 0 | 70 | 5 | 3 | 2 | 30466 |
|  | 0 | 0 | 0 | 0 | 0 | 99.7 | 0 | 0 | 0 | 0.01 | 0.01 | 0.01 | 0.01 | 0 | 0.23 | 0.02 | 0.01 | 0.01 |  |
| **10** | 0 | 0 | 0 | 0 | 0 | 1 | 0 | 0 | 0 | 1292 | 0 | 0 | 0 | 1 | 0 | 0 | 0 | 0 | 1294 |
|  | 0 | 0 | 0 | 0 | 0 | 0.08 | 0 | 0 | 0 | 99.85 | 0 | 0 | 0 | 0.08 | 0 | 0 | 0 | 0 |  |
| **11** | 17 | 0 | 0 | 0 | 0 | 0 | 0 | 0 | 0 | 0 | 0 | 0 | 0 | 699 | 0 | 0 | 0 | 0 | 716 |
|  | 2.37 | 0 | 0 | 0 | 0 | 0 | 0 | 0 | 0 | 0 | 0 | 0 | 0 | 97.63 | 0 | 0 | 0 | 0 |  |
| **13** | 0 | 0 | 0 | 0 | 0 | 0 | 0 | 0 | 0 | 0 | 0 | 0 | 0 | 0 | 582 | 0 | 0 | 0 | 582 |
|  | 0 | 0 | 0 | 0 | 0 | 0 | 0 | 0 | 0 | 0 | 0 | 0 | 0 | 0 | 100 | 0 | 0 | 0 |  |
| **2** | 0 | 0 | 1 | 0 | 0 | 0 | 0 | 434 | 0 | 4 | 0 | 186 | 5 | 0 | 0 | 0 | 0 | 0 | 630 |
|  | 0 | 0 | 0.16 | 0 | 0 | 0 | 0 | 68.89 | 0 | 0.63 | 0 | 29.52 | 0.79 | 0 | 0 | 0 | 0 | 0 |  |
| **3** | 2 | 0 | 0 | 1 | 0 | 0 | 29276 | 1 | 19 | 0 | 0 | 0 | 0 | 112 | 173 | 12 | 0 | 0 | 29596 |
|  | 0.01 | 0 | 0 | 0 | 0 | 0 | 98.92 | 0 | 0.06 | 0 | 0 | 0 | 0 | 0.38 | 0.58 | 0.04 | 0 | 0 |  |
| **4** | 0 | 3 | 0 | 0 | 0 | 0 | 0 | 7 | 0 | 0 | 0 | 1933 | 3 | 0 | 0 | 1 | 0 | 0 | 1947 |
|  | 0 | 0.15 | 0 | 0 | 0 | 0 | 0 | 0.36 | 0 | 0 | 0 | 99.28 | 0.15 | 0 | 0 | 0.05 | 0 | 0 |  |
| **5** | 5985 | 0 | 0 | 0 | 0 | 0 | 1 | 0 | 0 | 0 | 0 | 0 | 0 | 1 | 10 | 5 | 0 | 0 | 6002 |
|  | 99.72 | 0 | 0 | 0 | 0 | 0 | 0.02 | 0 | 0 | 0 | 0 | 0 | 0 | 0.02 | 0.17 | 0.08 | 0 | 0 |  |
| **6** | 1 | 0 | 0 | 0 | 0 | 0 | 0 | 0 | 0 | 0 | 0 | 0 | 0 | 1816 | 0 | 0 | 0 | 0 | 1817 |
|  | 0.06 | 0 | 0 | 0 | 0 | 0 | 0 | 0 | 0 | 0 | 0 | 0 | 0 | 99.94 | 0 | 0 | 0 | 0 |  |
| **7** | 0 | 0 | 0 | 0 | 1 | 0 | 2 | 0 | 0 | 0 | 0 | 0 | 0 | 0 | 1 | 2804 | 0 | 0 | 2808 |
|  | 0 | 0 | 0 | 0 | 0.04 | 0 | 0.07 | 0 | 0 | 0 | 0 | 0 | 0 | 0 | 0.04 | 99.86 | 0 | 0 |  |
| **9** | 4 | 0 | 12 | 0 | 0 | 0 | 0 | 1 | 0 | 1 | 0 | 0 | 0 | 0 | 7169 | 4 | 0 | 0 | 7191 |
|  | 0.06 | 0 | 0.17 | 0 | 0 | 0 | 0 | 0.01 | 0 | 0.01 | 0 | 0 | 0 | 0 | 99.69 | 0.06 | 0 | 0 |  |
| **99** | 0 | 0 | 0 | 0 | 0 | 0 | 0 | 0 | 38 | 0 | 0 | 485 | 0 | 2 | 8 | 290 | 0 | 0 | 823 |
|  | 0 | 0 | 0 | 0 | 0 | 0 | 0 | 0 | 4.62 | 0 | 0 | 58.93 | 0 | 0.24 | 0.97 | 35.24 | 0 | 0 |  |
| **Total** | 6009 | 3 | 13 | 1 | 1 | 30377 | 29280 | 443 | 57 | 1299 | 2 | 2606 | 11 | 2631 | 8013 | 3121 | 3 | 2 | 83872 |
|  | 7.16 | 0 | 0.02 | 0 | 0 | 36.22 | 34.91 | 0.53 | 0.07 | 1.55 | 0 | 3.11 | 0.01 | 3.14 | 9.55 | 3.72 | 0 | 0 | 100 |

**Supplementary Table 3s.**
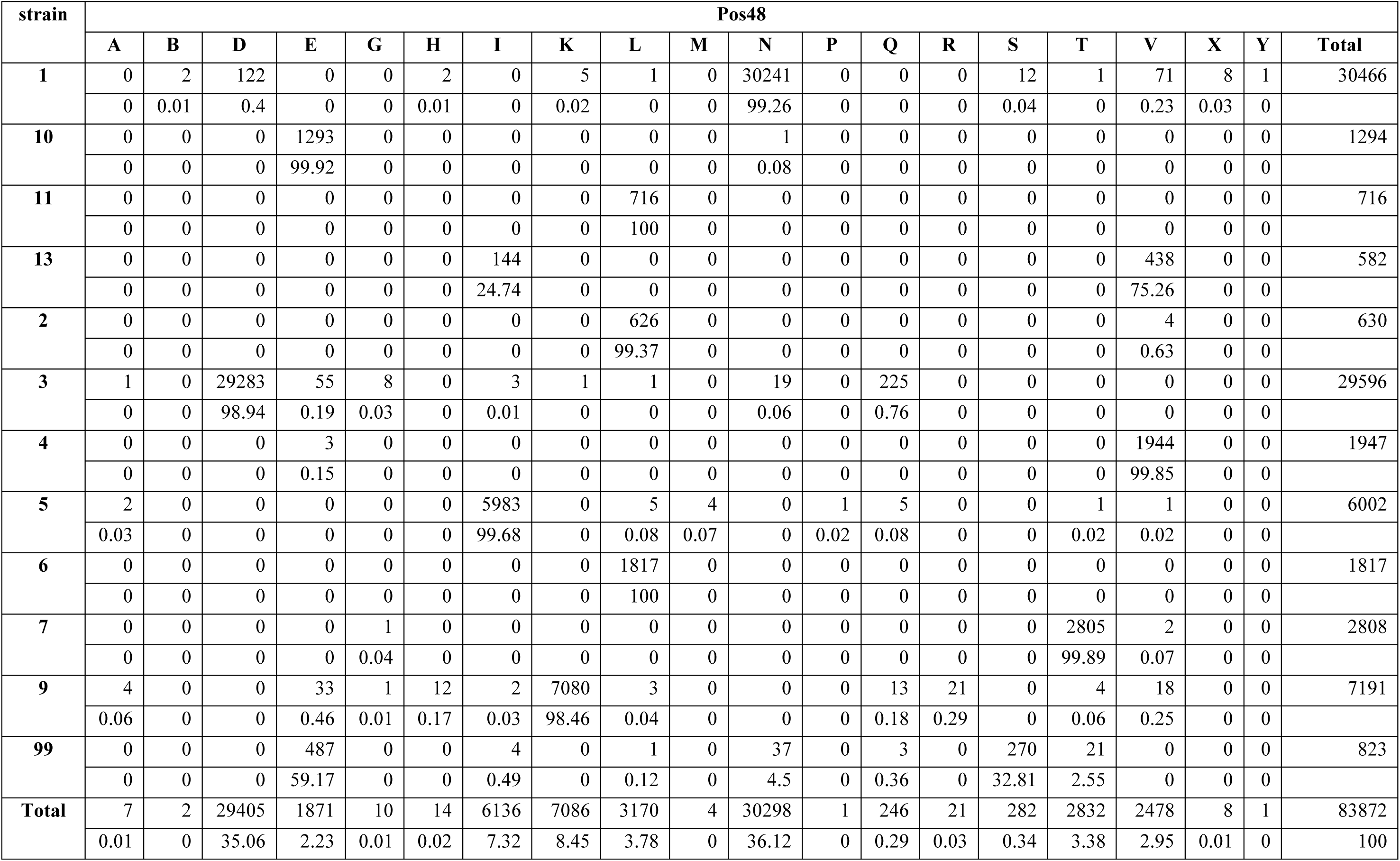
The distribution of amino acids in HA Position 48 among different HA types.

**Supplementary Table 4s.** The distribution of amino acids in NA Position 111 among different NA types.

| strain | Pos111 |  |  |  |  |  |  |  |  |  |  |  |  |  |  |  |  |  |  |  |  |  |  |
| --- | --- | --- | --- | --- | --- | --- | --- | --- | --- | --- | --- | --- | --- | --- | --- | --- | --- | --- | --- | --- | --- | --- | --- |
|  | - | A | C | D | E | F | G | H | I | K | L | M | N | P | Q | R | S | T | V | W | X | Y | Total |
| **1** | 0 | 1 | 4 | 0 | 2 | 107 | 26 | 40 | 6 | 24574 | 26 | 0 | 5 | 1 | 6 | 167 | 62 | 2993 | 5 | 0 | 6 | 0 | 28031 |
|  | 0 | 0 | 0.01 | 0 | 0.01 | 0.38 | 0.09 | 0.14 | 0.02 | 87.67 | 0.09 | 0 | 0.02 | 0 | 0.02 | 0.6 | 0.22 | 10.68 | 0.02 | 0 | 0.02 | 0 |  |
| **2** | 8 | 101 | 23 | 27 | 2 | 3 | 31282 | 0 | 1161 | 1 | 3 | 0 | 18 | 0 | 135 | 2 | 472 | 2 | 13 | 2 | 3 | 71 | 33329 |
|  | 0.02 | 0.3 | 0.07 | 0.08 | 0.01 | 0.01 | 93.86 | 0 | 3.48 | 0 | 0.01 | 0 | 0.05 | 0 | 0.41 | 0.01 | 1.42 | 0.01 | 0.04 | 0.01 | 0.01 | 0.21 |  |
| **3** | 0 | 6 | 0 | 3 | 0 | 0 | 1 | 1 | 0 | 15 | 33 | 0 | 1613 | 0 | 0 | 0 | 2 | 6 | 0 | 0 | 0 | 0 | 1680 |
|  | 0 | 0.36 | 0 | 0.18 | 0 | 0 | 0.06 | 0.06 | 0 | 0.89 | 1.96 | 0 | 96.01 | 0 | 0 | 0 | 0.12 | 0.36 | 0 | 0 | 0 | 0 |  |
| **4** | 0 | 0 | 0 | 0 | 0 | 0 | 375 | 0 | 0 | 0 | 0 | 0 | 0 | 0 | 0 | 0 | 0 | 0 | 0 | 0 | 0 | 0 | 375 |
|  | 0 | 0 | 0 | 0 | 0 | 0 | 100 | 0 | 0 | 0 | 0 | 0 | 0 | 0 | 0 | 0 | 0 | 0 | 0 | 0 | 0 | 0 |  |
| **5** | 0 | 0 | 0 | 0 | 6 | 141 | 0 | 0 | 0 | 0 | 0 | 0 | 0 | 0 | 0 | 0 | 0 | 0 | 434 | 0 | 0 | 0 | 581 |
|  | 0 | 0 | 0 | 0 | 1.03 | 24.27 | 0 | 0 | 0 | 0 | 0 | 0 | 0 | 0 | 0 | 0 | 0 | 0 | 74.7 | 0 | 0 | 0 |  |
| **6** | 0 | 176 | 0 | 2076 | 23 | 0 | 23 | 1 | 0 | 0 | 355 | 0 | 171 | 0 | 8 | 0 | 7 | 4 | 0 | 0 | 0 | 3 | 2847 |
|  | 0 | 6.18 | 0 | 72.92 | 0.81 | 0 | 0.81 | 0.04 | 0 | 0 | 12.47 | 0 | 6.01 | 0 | 0.28 | 0 | 0.25 | 0.14 | 0 | 0 | 0 | 0.11 |  |
| **7** | 0 | 0 | 0 | 0 | 315 | 0 | 3 | 2 | 15 | 0 | 0 | 1 | 0 | 0 | 714 | 0 | 0 | 2 | 0 | 0 | 0 | 47 | 1099 |
|  | 0 | 0 | 0 | 0 | 28.66 | 0 | 0.27 | 0.18 | 1.36 | 0 | 0 | 0.09 | 0 | 0 | 64.97 | 0 | 0 | 0.18 | 0 | 0 | 0 | 4.28 |  |
| **8** | 0 | 0 | 0 | 0 | 0 | 0 | 0 | 2787 | 0 | 0 | 3 | 0 | 0 | 0 | 0 | 0 | 0 | 0 | 1 | 0 | 1 | 0 | 2792 |
|  | 0 | 0 | 0 | 0 | 0 | 0 | 0 | 99.82 | 0 | 0 | 0.11 | 0 | 0 | 0 | 0 | 0 | 0 | 0 | 0.04 | 0 | 0.04 | 0 |  |
| **9** | 0 | 0 | 0 | 0 | 844 | 0 | 2 | 0 | 0 | 1 | 824 | 0 | 0 | 0 | 1 | 1 | 0 | 0 | 0 | 0 | 0 | 0 | 1673 |
|  | 0 | 0 | 0 | 0 | 50.45 | 0 | 0.12 | 0 | 0 | 0.06 | 49.25 | 0 | 0 | 0 | 0.06 | 0.06 | 0 | 0 | 0 | 0 | 0 | 0 |  |
| **Total** | 8 | 284 | 27 | 2106 | 1192 | 251 | 31712 | 2831 | 1182 | 24591 | 1244 | 1 | 1807 | 1 | 864 | 170 | 543 | 3007 | 453 | 2 | 10 | 121 | 72407 |
|  | 0.01 | 0.39 | 0.04 | 2.91 | 1.65 | 0.35 | 43.8 | 3.91 | 1.63 | 33.96 | 1.72 | 0 | 2.5 | 0 | 1.19 | 0.23 | 0.75 | 4.15 | 0.63 | 0 | 0.01 | 0.17 | 100 |

**Supplementary Table 5s.** The distribution of amino acids in NA Position 409 among different NA types.

| strain | Pos409 |  |  |  |  |  |  |  |  |  |  |  |  |  |  |  |  |  |  |  |  |  |  |
| --- | --- | --- | --- | --- | --- | --- | --- | --- | --- | --- | --- | --- | --- | --- | --- | --- | --- | --- | --- | --- | --- | --- | --- |
|  | - | A | C | D | E | F | G | H | I | K | L | M | N | P | Q | R | S | T | V | W | X | Y | Total |
| 1 | 0 | 1 | 11 | 0 | 20 | 4 | 2996 | 24656 | 3 | 7 | 5 | 0 | 2 | 126 | 4 | 101 | 2 | 0 | 40 | 50 | 2 | 1 | 28031 |
|  | 0 | 0 | 0.04 | 0 | 0.07 | 0.01 | 10.69 | 87.96 | 0.01 | 0.02 | 0.02 | 0 | 0.01 | 0.45 | 0.01 | 0.36 | 0.01 | 0 | 0.14 | 0.18 | 0.01 | 0 |  |
| 2 | 20 | 0 | 2 | 1 | 275 | 57 | 216 | 0 | 30992 | 6 | 23 | 14 | 43 | 5 | 1 | 36 | 86 | 7 | 1542 | 1 | 2 | 0 | 33329 |
|  | 0.06 | 0 | 0.01 | 0 | 0.83 | 0.17 | 0.65 | 0 | 92.99 | 0.02 | 0.07 | 0.04 | 0.13 | 0.02 | 0 | 0.11 | 0.26 | 0.02 | 4.63 | 0 | 0.01 | 0 |  |
| 3 | 0 | 0 | 0 | 1 | 2 | 1627 | 0 | 1 | 14 | 25 | 1 | 0 | 4 | 0 | 0 | 0 | 3 | 0 | 2 | 0 | 0 | 0 | 1680 |
|  | 0 | 0 | 0 | 0.06 | 0.12 | 96.85 | 0 | 0.06 | 0.83 | 1.49 | 0.06 | 0 | 0.24 | 0 | 0 | 0 | 0.18 | 0 | 0.12 | 0 | 0 | 0 |  |
| 4 | 0 | 0 | 0 | 0 | 0 | 0 | 0 | 0 | 375 | 0 | 0 | 0 | 0 | 0 | 0 | 0 | 0 | 0 | 0 | 0 | 0 | 0 | 375 |
|  | 0 | 0 | 0 | 0 | 0 | 0 | 0 | 0 | 100 | 0 | 0 | 0 | 0 | 0 | 0 | 0 | 0 | 0 | 0 | 0 | 0 | 0 |  |
| 5 | 0 | 0 | 0 | 0 | 0 | 0 | 0 | 0 | 142 | 0 | 0 | 6 | 0 | 0 | 0 | 0 | 0 | 432 | 1 | 0 | 0 | 0 | 581 |
|  | 0 | 0 | 0 | 0 | 0 | 0 | 0 | 0 | 24.44 | 0 | 0 | 1.03 | 0 | 0 | 0 | 0 | 0 | 74.35 | 0.17 | 0 | 0 | 0 |  |
| 6 | 0 | 2228 | 3 | 0 | 10 | 530 | 9 | 0 | 0 | 8 | 0 | 0 | 1 | 2 | 0 | 1 | 35 | 16 | 4 | 0 | 0 | 0 | 2847 |
|  | 0 | 78.26 | 0.11 | 0 | 0.35 | 18.62 | 0.32 | 0 | 0 | 0.28 | 0 | 0 | 0.04 | 0.07 | 0 | 0.04 | 1.23 | 0.56 | 0.14 | 0 | 0 | 0 |  |
| 7 | 0 | 0 | 0 | 15 | 0 | 315 | 0 | 0 | 761 | 0 | 1 | 0 | 3 | 0 | 0 | 2 | 2 | 0 | 0 | 0 | 0 | 0 | 1099 |
|  | 0 | 0 | 0 | 1.36 | 0 | 28.66 | 0 | 0 | 69.24 | 0 | 0.09 | 0 | 0.27 | 0 | 0 | 0.18 | 0.18 | 0 | 0 | 0 | 0 | 0 |  |
| 8 | 0 | 1 | 0 | 0 | 1 | 0 | 0 | 0 | 0 | 1 | 0 | 0 | 0 | 2788 | 0 | 0 | 0 | 0 | 1 | 0 | 0 | 0 | 2792 |
|  | 0 | 0.04 | 0 | 0 | 0.04 | 0 | 0 | 0 | 0 | 0.04 | 0 | 0 | 0 | 99.86 | 0 | 0 | 0 | 0 | 0.04 | 0 | 0 | 0 |  |
| 9 | 0 | 0 | 0 | 0 | 0 | 1 | 3 | 0 | 0 | 0 | 0 | 0 | 0 | 0 | 0 | 0 | 845 | 0 | 0 | 824 | 0 | 0 | 1673 |
|  | 0 | 0 | 0 | 0 | 0 | 0.06 | 0.18 | 0 | 0 | 0 | 0 | 0 | 0 | 0 | 0 | 0 | 50.51 | 0 | 0 | 49.25 | 0 | 0 |  |
| Total | 20 | 2230 | 16 | 17 | 308 | 2534 | 3224 | 24657 | 32287 | 47 | 30 | 20 | 53 | 2921 | 5 | 140 | 973 | 455 | 1590 | 875 | 4 | 1 | 72407 |
|  | 0.03 | 3.08 | 0.02 | 0.02 | 0.43 | 3.5 | 4.45 | 34.05 | 44.59 | 0.06 | 0.04 | 0.03 | 0.07 | 4.03 | 0.01 | 0.19 | 1.34 | 0.63 | 2.2 | 1.21 | 0.01 | 0 | 100 |

**Supplementary Table 6s.** The distribution of amino acids in NA Position 34 among different NA types.

| Table 6s. The distribution of amino acids in NA Position 34 among different NA types. |  |  |  |  |  |  |  |  |  |  |  |  |  |  |  |  |  |  |  |  |  |
| --- | --- | --- | --- | --- | --- | --- | --- | --- | --- | --- | --- | --- | --- | --- | --- | --- | --- | --- | --- | --- | --- |
| strain | Pos34 |  |  |  |  |  |  |  |  |  |  |  |  |  |  |  |  |  |  |  |  |
|  | A | C | D | E | F | G | H | I | K | L | M | N | P | Q | R | S | T | V | W | X | Total |
| 1 | 2226 | 0 | 0 | 0 | 3 | 10 | 0 | 13327 | 2 | 20 | 49 | 21 | 2 | 1 | 0 | 7 | 22 | 12337 | 3 | 1 | 28031 |
|  | 7.94 | 0 | 0 | 0 | 0.01 | 0.04 | 0 | 47.54 | 0.01 | 0.07 | 0.17 | 0.07 | 0.01 | 0 | 0 | 0.02 | 0.08 | 44.01 | 0.01 | 0 |  |
| 2 | 29 | 2 | 1 | 0 | 0 | 16 | 1 | 18 | 167 | 2 | 6 | 2 | 1 | 0 | 2 | 5 | 33075 | 0 | 0 | 2 | 33329 |
|  | 0.09 | 0.01 | 0 | 0 | 0 | 0.05 | 0 | 0.05 | 0.5 | 0.01 | 0.02 | 0.01 | 0 | 0 | 0.01 | 0.02 | 99.24 | 0 | 0 | 0.01 |  |
| 3 | 1 | 0 | 0 | 0 | 2 | 0 | 0 | 1 | 0 | 0 | 0 | 0 | 0 | 0 | 0 | 0 | 0 | 1676 | 0 | 0 | 1680 |
|  | 0.06 | 0 | 0 | 0 | 0.12 | 0 | 0 | 0.06 | 0 | 0 | 0 | 0 | 0 | 0 | 0 | 0 | 0 | 99.76 | 0 | 0 |  |
| 4 | 0 | 0 | 0 | 0 | 375 | 0 | 0 | 0 | 0 | 0 | 0 | 0 | 0 | 0 | 0 | 0 | 0 | 0 | 0 | 0 | 375 |
|  | 0 | 0 | 0 | 0 | 100 | 0 | 0 | 0 | 0 | 0 | 0 | 0 | 0 | 0 | 0 | 0 | 0 | 0 | 0 | 0 |  |
| 5 | 0 | 0 | 0 | 0 | 0 | 0 | 0 | 569 | 0 | 0 | 1 | 0 | 0 | 0 | 0 | 0 | 0 | 11 | 0 | 0 | 581 |
|  | 0 | 0 | 0 | 0 | 0 | 0 | 0 | 97.93 | 0 | 0 | 0.17 | 0 | 0 | 0 | 0 | 0 | 0 | 1.89 | 0 | 0 |  |
| 6 | 0 | 0 | 0 | 1 | 0 | 2835 | 0 | 8 | 0 | 0 | 0 | 0 | 0 | 0 | 2 | 0 | 0 | 0 | 0 | 1 | 2847 |
|  | 0 | 0 | 0 | 0.04 | 0 | 99.58 | 0 | 0.28 | 0 | 0 | 0 | 0 | 0 | 0 | 0.07 | 0 | 0 | 0 | 0 | 0.04 |  |
| 7 | 0 | 0 | 0 | 0 | 0 | 0 | 0 | 0 | 0 | 0 | 0 | 0 | 0 | 0 | 0 | 1099 | 0 | 0 | 0 | 0 | 1099 |
|  | 0 | 0 | 0 | 0 | 0 | 0 | 0 | 0 | 0 | 0 | 0 | 0 | 0 | 0 | 0 | 100 | 0 | 0 | 0 | 0 |  |
| 8 | 0 | 0 | 0 | 0 | 2 | 0 | 0 | 5 | 0 | 2648 | 0 | 0 | 0 | 0 | 0 | 1 | 1 | 135 | 0 | 0 | 2792 |
|  | 0 | 0 | 0 | 0 | 0.07 | 0 | 0 | 0.18 | 0 | 94.84 | 0 | 0 | 0 | 0 | 0 | 0.04 | 0.04 | 4.84 | 0 | 0 |  |
| 9 | 0 | 0 | 0 | 1 | 0 | 1671 | 0 | 0 | 0 | 0 | 0 | 0 | 0 | 0 | 1 | 0 | 0 | 0 | 0 | 0 | 1673 |
|  | 0 | 0 | 0 | 0.06 | 0 | 99.88 | 0 | 0 | 0 | 0 | 0 | 0 | 0 | 0 | 0.06 | 0 | 0 | 0 | 0 | 0 |  |
| Total | 2256 | 2 | 1 | 2 | 382 | 4532 | 1 | 13928 | 169 | 2670 | 56 | 23 | 3 | 1 | 5 | 1112 | 33098 | 14159 | 3 | 4 | 72407 |
|  | 3.12 | 0 | 0 | 0 | 0.53 | 6.26 | 0 | 19.24 | 0.23 | 3.69 | 0.08 | 0.03 | 0 | 0 | 0.01 | 1.54 | 45.71 | 19.55 | 0 | 0.01 | 100 |

**Supplementary Table 7s.** The distribution of amino acids in NA Position 45 among different NA types.

| strain | Pos45 |  |  |  |  |  |  |  |  |  |  |  |  |  |  |  |  |  |  |  |  |  |
| --- | --- | --- | --- | --- | --- | --- | --- | --- | --- | --- | --- | --- | --- | --- | --- | --- | --- | --- | --- | --- | --- | --- |
|  | A | C | D | E | F | G | H | I | K | L | M | N | P | Q | R | S | T | V | W | X | Y | Total |
| 1 | 1 | 61 | 0 | 43 | 2 | 0 | 5110 | 4 | 158 | 19 | 0 | 1067 | 16 | 21112 | 76 | 23 | 2 | 1 | 3 | 3 | 330 | 28031 |
|  | 0 | 0.22 | 0 | 0.15 | 0.01 | 0 | 18.23 | 0.01 | 0.56 | 0.07 | 0 | 3.81 | 0.06 | 75.32 | 0.27 | 0.08 | 0.01 | 0 | 0.01 | 0.01 | 1.18 |  |
| 2 | 8 | 2 | 1 | 2 | 141 | 0 | 32 | 9 | 0 | 480 | 0 | 24 | 28303 | 21 | 8 | 4110 | 155 | 2 | 8 | 13 | 10 | 33329 |
|  | 0.02 | 0.01 | 0 | 0.01 | 0.42 | 0 | 0.1 | 0.03 | 0 | 1.44 | 0 | 0.07 | 84.92 | 0.06 | 0.02 | 12.33 | 0.47 | 0.01 | 0.02 | 0.04 | 0.03 |  |
| 3 | 6 | 0 | 1 | 126 | 0 | 138 | 1 | 2 | 0 | 0 | 4 | 0 | 0 | 0 | 0 | 0 | 11 | 1391 | 0 | 0 | 0 | 1680 |
|  | 0.36 | 0 | 0.06 | 7.5 | 0 | 8.21 | 0.06 | 0.12 | 0 | 0 | 0.24 | 0 | 0 | 0 | 0 | 0 | 0.65 | 82.8 | 0 | 0 | 0 |  |
| 4 | 0 | 0 | 0 | 363 | 0 | 12 | 0 | 0 | 0 | 0 | 0 | 0 | 0 | 0 | 0 | 0 | 0 | 0 | 0 | 0 | 0 | 375 |
|  | 0 | 0 | 0 | 96.8 | 0 | 3.2 | 0 | 0 | 0 | 0 | 0 | 0 | 0 | 0 | 0 | 0 | 0 | 0 | 0 | 0 | 0 |  |
| 5 | 0 | 147 | 0 | 0 | 2 | 0 | 0 | 0 | 0 | 0 | 0 | 1 | 186 | 0 | 0 | 59 | 186 | 0 | 0 | 0 | 0 | 581 |
|  | 0 | 25.3 | 0 | 0 | 0.34 | 0 | 0 | 0 | 0 | 0 | 0 | 0.17 | 32.01 | 0 | 0 | 10.15 | 32.01 | 0 | 0 | 0 | 0 |  |
| 6 | 35 | 0 | 0 | 0 | 0 | 0 | 0 | 627 | 0 | 6 | 188 | 9 | 53 | 0 | 0 | 189 | 1198 | 542 | 0 | 0 | 0 | 2847 |
|  | 1.23 | 0 | 0 | 0 | 0 | 0 | 0 | 22.02 | 0 | 0.21 | 6.6 | 0.32 | 1.86 | 0 | 0 | 6.64 | 42.08 | 19.04 | 0 | 0 | 0 |  |
| 7 | 3 | 0 | 46 | 322 | 0 | 0 | 0 | 0 | 617 | 0 | 0 | 8 | 2 | 0 | 101 | 0 | 0 | 0 | 0 | 0 | 0 | 1099 |
|  | 0.27 | 0 | 4.19 | 29.3 | 0 | 0 | 0 | 0 | 56.14 | 0 | 0 | 0.73 | 0.18 | 0 | 9.19 | 0 | 0 | 0 | 0 | 0 | 0 |  |
| 8 | 0 | 2775 | 0 | 0 | 0 | 0 | 0 | 0 | 0 | 0 | 0 | 0 | 0 | 0 | 0 | 1 | 0 | 0 | 1 | 0 | 15 | 2792 |
|  | 0 | 99.39 | 0 | 0 | 0 | 0 | 0 | 0 | 0 | 0 | 0 | 0 | 0 | 0 | 0 | 0.04 | 0 | 0 | 0.04 | 0 | 0.54 |  |
| 9 | 0 | 0 | 0 | 0 | 0 | 0 | 1494 | 0 | 0 | 0 | 0 | 64 | 2 | 26 | 81 | 6 | 0 | 0 | 0 | 0 | 0 | 1673 |
|  | 0 | 0 | 0 | 0 | 0 | 0 | 89.3 | 0 | 0 | 0 | 0 | 3.83 | 0.12 | 1.55 | 4.84 | 0.36 | 0 | 0 | 0 | 0 | 0 |  |
| Total | 53 | 2985 | 48 | 856 | 145 | 150 | 6637 | 642 | 775 | 505 | 192 | 1173 | 28562 | 21159 | 266 | 4388 | 1552 | 1936 | 12 | 16 | 355 | 72407 |
|  | 0.07 | 4.12 | 0.07 | 1.18 | 0.2 | 0.21 | 9.17 | 0.89 | 1.07 | 0.7 | 0.27 | 1.62 | 39.45 | 29.22 | 0.37 | 6.06 | 2.14 | 2.67 | 0.02 | 0.02 | 0.49 | 100 |

**Supplementary Table 8s.**
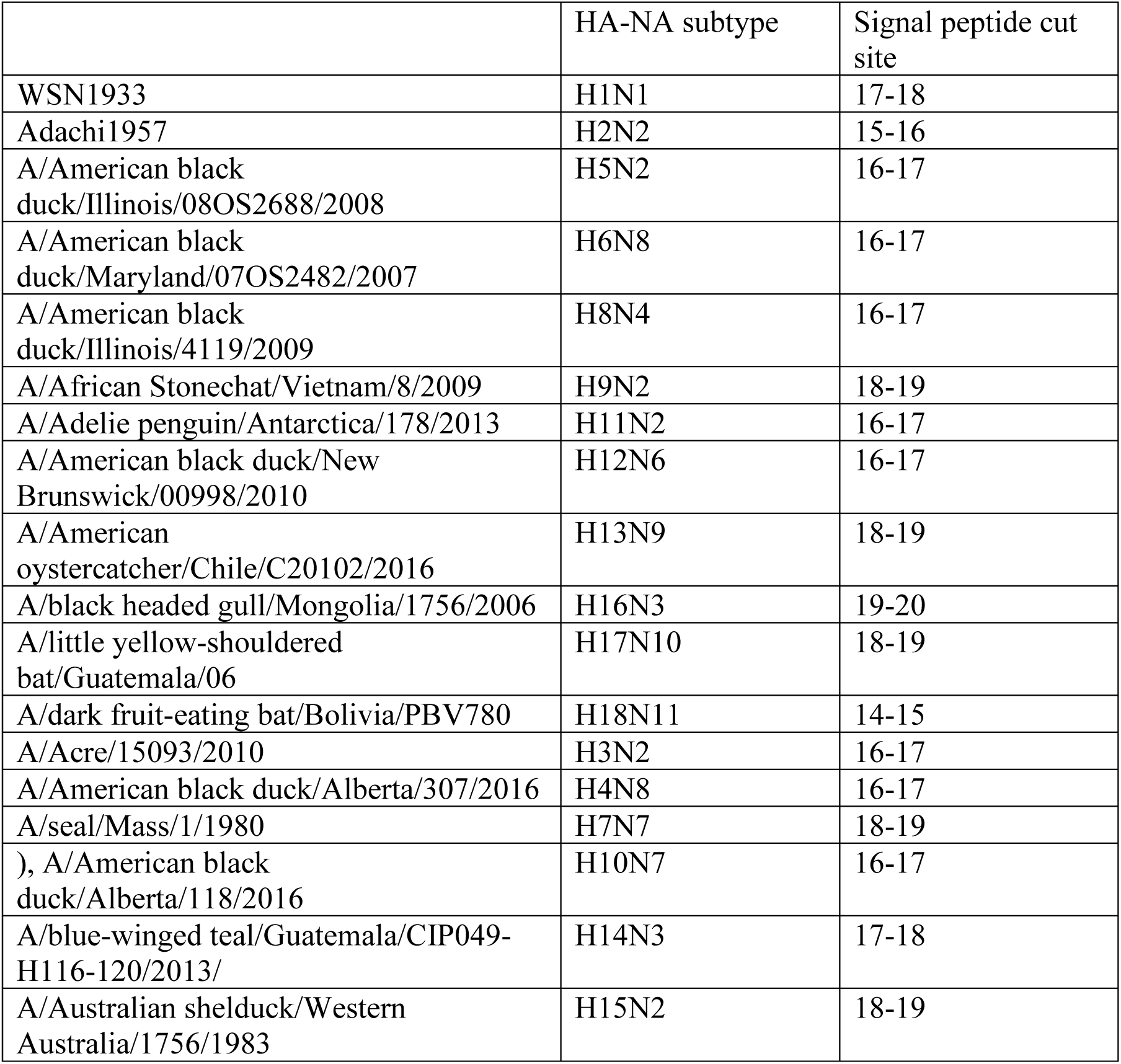
List of HA sequences used in the alignment.

**Supplementary Table 9s.** List of NA sequences used in the alignment.

**SupplementaryTable 9s List of NA sequences used in the alignment**
|  | HA-NA subtype | Signal peptide cut site |
| --- | --- | --- |
| A/swine/Iowa/A02432126/2019 | H1N1 | // |
| A/Northern Shoveler/California/D1614933/2016 | H8N4 | // |
| A/ruddy turnstone/Delaware Bay/592/2018 | H12N5 | // |
| A/ruddy turnstone/Delaware Bay/265/2018 | H3N8 | // |
| A/Florida/8584/2019 | H3N2 | // |
| A/Mallard/California/D1516363/2015 | H2N3 | // |
| A/Mallard/California/D1610190/2016 | H4N6 | // |
| A/red knot/Delaware Bay/393/201 | H11N7 | // |
| A/mallard/Alberta/149/2018 | H2N9 | // |

